# Long-read whole-genome sequencing-based concurrent haplotyping and aneuploidy profiling of single cells

**DOI:** 10.1101/2024.09.24.614469

**Authors:** Yan Zhao, Olga Tsuiko, Tatjana Jatsenko, Greet Peeters, Erika Souche, Mathilde Geysens, Eftychia Dimitriadou, Arne Vanhie, Karen Peeraer, Sophie Debrock, Hilde Van Esch, Joris Robert Vermeesch

## Abstract

Long-read whole-genome sequencing (lrWGS) enhances haplotyping by providing more phasing information per read compared to short-read sequencing. However, its use for single-cell haplotype phasing remains underexplored. This proof-of-concept study examines lrWGS data from single cells for small variant (SNV and indel) calling and haplotyping using the Genome in a Bottle (GIAB) Ashkenazi trio. lrWGS was performed on single-cell (1 cell) and multi-cell (10 cells) samples from the offspring. Chromosome-length haplotypes were obtained by leveraging both long reads and pedigree information. These haplotypes were further refined by replacing them with matched parental haplotypes. In single-cell and multi-cell samples, 92% and 98% of heterozygous SNVs, and 74% and 78% of heterozygous indels were accurately haplotyped. Applied to human embryos for preimplantation genetic testing (PGT), lrWGS demonstrated 100% consistency with array-based methods for detecting monogenic disorders, without requiring phasing references. Aneuploidies were accurately detected, with insights into the mechanistic origins of chromosomal abnormalities inferred from the parental unique allele fractions. We show that lrWGS-based concurrent haplotyping and aneuploidy profiling of single cells provides an alternative to current PGT methods, with applications potential in areas such as cell-based prenatal diagnosis and animal and plant breeding.

## Introduction

Most mammalian genomes are diploid, consisting of one haploid set of chromosomes from each parent. Haplotyping reconstructs the unique nucleotide content of the two homologous chromosome sets, known as haplotypes. This process is crucial because the haplotypes can have different functional roles (1). Recent advancements in long-read sequencing technologies provide read lengths over 10 kilobases and accuracy comparable to next-generation sequencing (NGS) (2). These improvements significantly enhance genome haplotyping, as individual long reads cover more heterozygous SNVs and provide haplotype information across extensive genomic regions, surpassing the capabilities of traditional SNP arrays and short-read data. The main long-read sequencing technologies are the single molecule real-time (SMRT) sequencing from Pacific Bioscience (PacBio) (3) and nanopore sequencing from Oxford Nanopore Technologies (ONT) (4). Studies have highlighted the effectiveness of long-read sequencing in variant identification, haplotyping, and genome assembly. Wenger et al. demonstrated the ability of ∼28x PacBio high-fidelity (HiFi) reads for high-performance small variant (SNV and indel) calling and phased 99.64% of called variants. (5). More recently, the Telomere-to-Telomere (T2T) Consortium created a complete human reference genome, T2T-CHM13, using PacBio HiFi reads and ONT ultralong reads (6).

In addition to enabling haplotyping and genome assembly for bulk samples, long-read sequencing also holds potential for facilitating haplotyping at the single-cell level. Genetic analysis for single cells is challenging due to the necessity of whole genome amplification (WGA), typically using techniques like multiple displacement amplification (MDA). WGA can introduce technical errors, including allele dropout (ADO) due to the failure to amplify one allele, false-positive errors resulting from polymerase infidelity, and coverage nonuniformity caused by uneven amplification(7). Due to WGA artifacts, haplotype-based analysis of single cells is crucial for areas like preimplantation genetic testing for monogenic disorders (PGT-M). In this context, embryos from couples at risk of transmitting genetic disorders to their offspring are tested for the inheritance of disease alleles using DNA from trophectoderm (TE) biopsies containing 5 to 10 cells or from single blastomere biopsies. Several genome-wide haplotyping methods for single cells have been developed, including karyomapping (8), siCHILD (9), One PGT (10) Haploseek (11), and scGBS (12). These methods utilize SNP arrays or NGS data for genotyping, which provide minimal or no haplotype information. As a result, genetic phasing is applied, requiring DNA samples from prospective parents and first-degree relative(s) (Carvalho *et al.*, 2020), which are not always available. Furthermore, in cases of de novo mutations (DNMs) in prospective parents, the variant loci cannot be phased through genetic phasing. Long-read data has the potential to directly phase both parents and embryo biopsies, including DNMs in the parents, without requiring relatives. Initial explorations focused on targeted long-read sequencing. For instance, Wu et al. conducted haplotype linkage analysis for the HBB gene by phasing the parents and TE biopsies using SMRT reads (13). Similarly, Tsuiko et al. explored both SMRT and ONT data in preclinical workup to infer the parental origin of DNMs (14). More recently, long-read whole-genome sequencing (lrWGS) has been employed for phasing parental genomes. Zhang et al. utilized ∼30x PacBio long-read data for phasing the parents and conducted reference-free PGT-M for three monogenetic diseases (15). However, the effectiveness of lrWGS for phasing single cells and its application in generic PGT remains unexplored. Hård et al. assessed variant calling and genome assembly with lrWGS data from single cells (16), but the limited sequencing depth of ∼ 5x HiFi reads constrains a thorough exploration of its potential for clinical applications.

Beyond haplotyping, SNP arrays and NGS-based single-cell haplotyping methods enable concurrent haplotype-aware aneuploidy profiling, facilitating the identification of chromosomal abnormalities in single cells and determining their mechanistic origins. This has significant clinical implications, as chromosomal abnormalities can arise during human gametogenesis and are common in early embryogenesis (17, 18). PGT for aneuploidy (PGT-A) prevents the transmission of chromosomally abnormal embryos and enhances the in vitro fertilization (IVF) success rate (19). Long-read sequencing has proven valuable for detecting aneuploidies (20) and segmental imbalances (21) in embryo biopsies. However, to our knowledge, the potential of long-read data to infer the mechanistic origins of aneuploidies in embryo biopsies has not yet been explored.

Here, we present the first comprehensive analysis of lrWGS data from human single cells at an adequate depth of ∼24x for SNV and indel calling, as well as haplotyping. Using a Genome in a Bottle (GIAB) trio consisting of HG002 (offspring), HG003 (father), and HG004 (mother) for benchmarking, we demonstrate the feasibility of lrWGS data for concurrent haplotyping and aneuploidy profiling of single cells without requiring additional phasing references. The clinical proof-of-concept application was validated in two PGT families, achieving 100% diagnostic concordance with SNP array-based PGT results (**Figure 1)**. This lrWGS-based PGT approach surpasses current methods with reduced clinical work-up, fewer family members involved, and a more comprehensive genomic analysis that integrates direct variant detection, haplotyping, and aneuploidy assessment. Furthermore, our data analysis strategy for concurrent haplotyping and aneuploidy profiling of single cells can be applied to other areas of single-cell genome analysis, such as cell-based prenatal diagnosis and animal and plant breeding.

**Figure 1.**
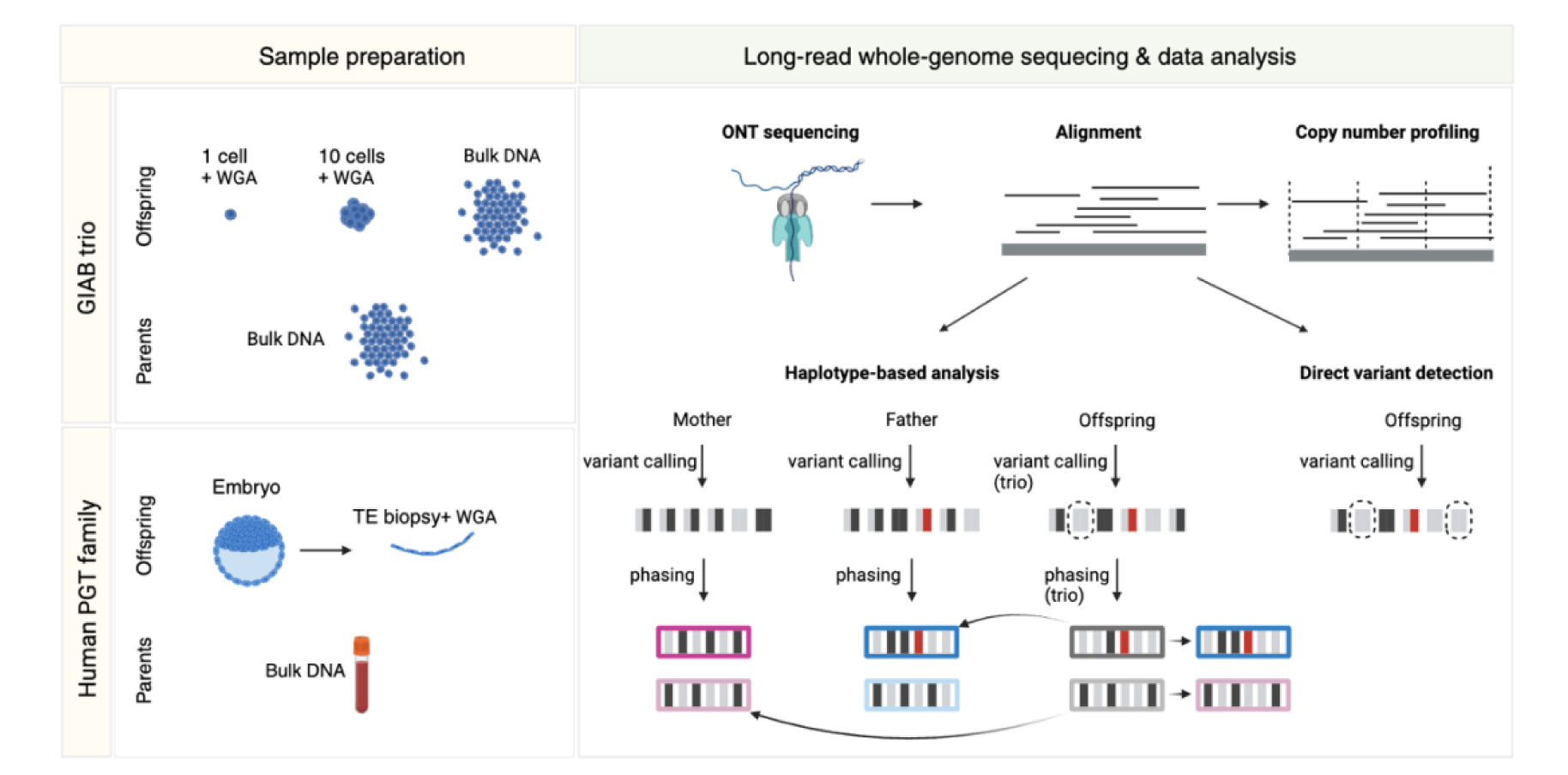
lrWGS-based concurrent haplotyping and aneuploidy profiling workflow. The benchmarking study utilized a Genome in a Bottle (GIAB) trio, consisting of HG002 (offspring), HG003 (father), and HG004 (mother). Single-cell (1 cell), multi-cell (10 cells), and bulk samples were collected from the HG002 cell line. The single-cell and multi-cell samples underwent WGA before lrWGS using nanopore technology. The long reads were then aligned to the human reference genome. Publicly available aligned lrWGS data for the parents were used in subsequent analyses. After alignment, aneuploidy profiling was performed on each sample. Two approaches were applied to determine whether a disease allele present in the parents was inherited by the offspring: haplotype-based analysis and direct variant detection. In the haplotype-based analysis, the parents underwent individual variant calling and phasing, while the offspring was analyzed in a trio setting, which incorporates parental information for better results. By comparing the offspring’s haplotypes with those of the parents, the inherited parental haplotypes were inferred, which were subsequently used for diagnosis. For direct variant detection, the offspring underwent independent variant calling without parental information, and the results were utilized for diagnosis. In variant calling results, different alleles are depicted as black or grey rectangles, with the “disease allele” highlighted in red. Each pair of rectangles represents a genotype, and those affected by ADO are indicated by dashed circles. After phasing, the two alleles in a genotype were organized into distinct haplotypes. In this example, the disease allele inherited by the offspring was identified by both haplotype-based analysis and direct variant detection. The utility of lrWGS-based haplotyping and aneuploidy profiling was further evaluated in families undergoing PGT-M, where TE biopsies from embryos and bulk DNA from the prospective parents were analyzed.

## Materials and methods

### Long-read whole-genome sequencing for the GIAB trio

Lymphoblastoid cell line of the offspring (HG002) from the GIAB Ashkenazi trio, consisting of HG002 (offspring), HG003 (father), and HG004 (mother) (Coriell Institute), was cultured in DMEM/F12, complemented with 10% FBS (Gibco). Single-cell (1 cell) and multi-cell (10 cells) samples were manually collected into 0.2ml tubes and DNA from these samples was amplified by MDA using the REPLI-g single cell kit (QIAGEN). ONT libraries were prepared on 3 µg of amplified material using the SQK-LSK114 kit, following ONT’s recommendations for WGA library preparations. A bulk sample was collected in parallel, and DNA was extracted using the genomic DNA purification kit (Monarch). Libraries were then prepared on 3 µg of bulk DNA using the SQK-LSK114 kit. All libraries were loaded (30 fmol) on the PromethION device for sequencing with R10.4.1 flow cells. lrWGS datasets for the father (HG003) and mother (HG004) were obtained from the Oxford Nanopore Open Datasets. Specifically, BAM files were downloaded from the AWS storage bucket at s3://ont-open-data/giab_lsk114_2022.12/ and downsampled to ∼ 24x, which is comparable to that of the offspring, using samtools (v 1.9) (22). Details regarding the base calling model and base caller used for each cell line sample are provided in **Supplementary Table 1**.

### Long-read whole-genome sequencing for human PGT families

This study was approved by the Ethical Committee of UZ/KU Leuven (S68291). The parents consented to the use of residuary Human Bodily Material (HBM) for scientific research at the start of their IVF treatment. IVF and universal PGT-related procedures have been performed according to the standard operating procedures (SOPs) of UZ Leuven. Specifically, for universal PGT, parental bulk DNA was extracted from whole blood. TE biopsy was performed on blastocyst-stage embryos, obtaining an average of five TE cells, which were subjected to WGA using the MDA method with the REPLI-g SC kit (Qiagen, Germany). Residual DNA from both parents and corresponding embryo biopsies (MDA amplified) is available at the hospital, and an aliquot of excess DNA material were utilized for this study. ONT library preparation and sequencing were done using the same procedures as for cell line samples, with the base calling model and base caller applied for each sample listed in **Supplementary Table 1**.

### Read preprocessing and mapping

Read quality was assessed using NanoPlot (v1.39.0) (23) and FastQC (v0.11.7). Reads with an average quality score below 9 or a length shorter than 500 bp were filtered out using NanoFilt (v2.8.0) (23). Subsequently, the processed reads were aligned to human reference genome hg38 using minimap2 (v2.12) (24). Mapping statistics were generated using samtools flagstat (v1.9) (22). Chimeric read count was determined by counting the number of unique reads for all alignments with 0x800 SAM flag. Depth of coverage for each genomic position was computed using samtools depth with -aa flag (v1.9) (22), and average depth of coverage was determined by dividing the sum of depths across all positions by the size of the genome.

### Variant calling

Clair3 (v0.1) (25) was used for variant calling with BAM file of a single individual. Clair3-Trio (v0.3) (26) was used for trio variant calling with BAM files of the parents and the offspring. For both strategies, default parameters were used and SNVs and short indels (≤50 base pairs) were called. GIAB benchmark data (v4.2.1) was used to assess variant calling performance on high-confidence regions. VCF files containing high confident small variants (SNVs and short indels) on autosomes and corresponding BED files containing high confident regions were obtained for each trio member from https://ftp-trace.ncbi.nlm.nih.gov/giab/ftp/release/AshkenazimTrio/. RTG Tools (v3.12.1) (27) was used for benchmarking analysis with the vcfeval command.

To include supplementary alignments in variant calling (omitted by default), a customized shell script was employed to modify the SAM flags of the supplementary alignments (2048 and 2064) as normal flags (0 and 16, respectively). Subsequently, the modified BAM files were utilized for variant calling with Clair3 (v0.1).

### Haplotype phasing

The default phasing mode of WhatsHap (v1.0) (28), which uses data from a single individual, was employed to phase variants identified by Clair3. The pedigree phasing mode of WhatsHap (v1.0) (28), utilizing data from both parents and the offspring, was used to phase parental variants called by Clair3 and offspring’s variants called by Clair3-Trio. For both modes, default parameters were applied to phase only SNVs, while the – indels flag was added to phase both SNVs and indels. Only variants with GQ values higher than 2 in VCF files were retained for phasing. Preliminary conservative paternal|maternal phasing data from GIAB (available at https://ftp-trace.ncbi.nlm.nih.gov/giab/ftp/release/AshkenazimTrio/HG002_NA24385_son/NISTv4.2.1/GRCh38/SupplementaryFiles/HG002_GRCh38_1_22_v4.2.1_benchmark_phased_MHCassembly_StrandSeqANDTrio.vcf.gz) was used as a benchmark for evaluating phasing performance with WhatsHap Compare (v1.0)(28).

### Inference of inherited parental haplotypes

To deduce the parental haplotypes inherited by the offspring, haplotypes of the offspring obtained from the pedigree phasing mode were compared with parental haplotypes from the default phasing mode. Only biallelic loci were retained for comparison, excluding those with unphased genotypes in either parent or identical homozygous genotypes in both parents. Each chromosome was divided into 1 Mb consecutive segments, referred to as comparison units, and haplotype comparisons were performed within each segment. Since in the resulting VCF file from pedigree phasing mode, haplotype alleles of the offspring are given as paternal|maternal, for each comparison unit, we compared the first haplotype of the offspring to the two paternal haplotypes and the second haplotype of the offspring to the two maternal haplotypes. During comparison, loci with unphased genotypes or genotypes that violated Mendelian inheritance rules in the offspring were disregarded. The inherited parental haplotypes were identified as those exhibiting the highest number of matched SNVs and were used to replace the original haplotype information in the offspring. For each locus of interest, the offspring’s genotype was determined from these inherited parental haplotypes.

### Performance evaluation for haplotype linkage analysis and direct variant detection

To evaluate the performance of both haplotype linkage analysis and direct variant detection, familial high-confidence regions were first obtained by intersecting the high-confidence regions of each trio member using bedtools multiinter (v2.27.1) (29). The familial high-confidence regions were then intersected with protein-coding gene regions (extracted from the genome annotation file downloaded from https://ftp.ebi.ac.uk/pub/databases/gencode/Gencode_human/release_42/gencode.v42.annotation.gtf.gz) using bedtools intersect (v2.27.1) (29). Within the resulting intersected regions, loci with parental genotype combinations of heterozygous (0/1) and homozygous reference (0/0) were selected. The performances of the two methods were then assessed by evaluating whether the genotypes of the offspring at these loci could be accurately identified from the inferred parental haplotypes (haplotype linkage analysis) or from default variant calling results (direct variant detection).

### De novo mutation screening

Genome-wide DNM screening was restricted to biallelic SNV loci within familial high confidence gene regions as detailed above. DNMs were identified by comparing the genotypes of the trio members. A locus was classified as DNM if both parental genotypes were homozygous reference while the offspring showed a different genotype. Further detailed analysis of the identified DNMs were done with a customized R script.

### Aneuploidy profiling

Aneuploidy profiling was performed using Nano-GLADIATOR (v1.0) (30) with a window size of 1,000,000 base pairs.

To determine parental haplotype contributions across the genome, we selected SNV loci where the parents exhibited differing homozygous genotypes (homozygous reference (0/0) for one parent and homozygous alternate (1/1) for the other). Only loci with GQ > 2 and depth (DP) between 5 and 50 in the offspring were retained. For each locus, we computed the paternal and maternal allele fractions for the offspring. If a parent’s genotype was 1/1, the allele frequency (AF) value for the offspring was used as the allele fraction for that parent. Conversely, 1 minus AF was used as the allele fraction if the parent’s genotype was 0/0. Subsequently, we grouped loci within fixed bin sizes of 1 MB and calculated the average paternal and maternal allele fractions within each bin. Bins containing fewer than 30,000 loci were excluded. The calculated average paternal and maternal allele fraction values then underwent Circular Binary Segmentation using the R package PSCBS. Finally, we plotted the mean parental allele fraction values for each bin along with the segmentations across the genome. The parental haplotype fraction for each chromosome was determined through visual inspection of the plot, enabling the identification of the parental origin of any aneuploidies.

To determine the mitotic or meiotic origin of an aneuploidy with identified parental origin, we analyzed SNV loci that were heterozygous in the parent causing the aneuploidy and, in the offspring, but homozygous in the other parent. For each locus in the offspring, the unique allele fraction (UAF) was inferred from the AF value, where the unique allele refers to the distinct allele among the four parental alleles. Loci were grouped within fixed bins of 1 MB, and the average UAF was calculated for each bin. Bins with fewer than 30,000 loci were excluded from the analysis. The average UAF was subjected to Circular Binary Segmentation using the R package PSCBS. Finally, the average UAF for each bin, along with segmentations, was plotted across the genome. The mitotic or meiotic origin of the aneuploidy was then inferred through visual inspection of the plot.

## Results

### SNV and indel calling with long-read whole-genome sequencing data from single cells

To evaluate the potential of lrWGS-based haplotyping for single cells, we took one single-cell and one multi-cell (10 cells) sample from the offspring of the GIAB trio, mimicking single blastomere and TE biopsy, respectively. Additionally, a bulk sample was included for comparison (**Figure 1**). We obtained 22-24x lrWGS data for the bulk, multi-cell, and single-cell samples, covering 95%, 94%, and 88% of the human genome, respectively **(Supplementary Table 2 and Supplementary Table 3)**.

Given that SNVs and indels are the primary focus of most PGT-M cases and serve as genetic markers for haplotype construction, we first performed SNV and indel calling using lrWGS data. Variant calling performance was assessed by comparing with GIAB benchmark data. Across all sample types, we observed better variant calling performance for SNVs than for indels **(Figure 2A)**. Among different sample types, single-cell data showed the lowest performance, while multi-cell data was more similar to bulk data. Specifically, multi-cell data exhibited an SNV F-measure of 0.9605, comparable to bulk data at 0.9925, whereas the F-measure for single-cell data decreased significantly to 0.6305 **(Figure 2A)**. Not surprisingly, for single-cell data, the sensitivity for heterozygous SNVs was notably lower (0.3844) compared to homozygous SNVs (0.8422) (**Supplementary Figure 1**), suggesting a high rate of ADO.

**Figure 2.**
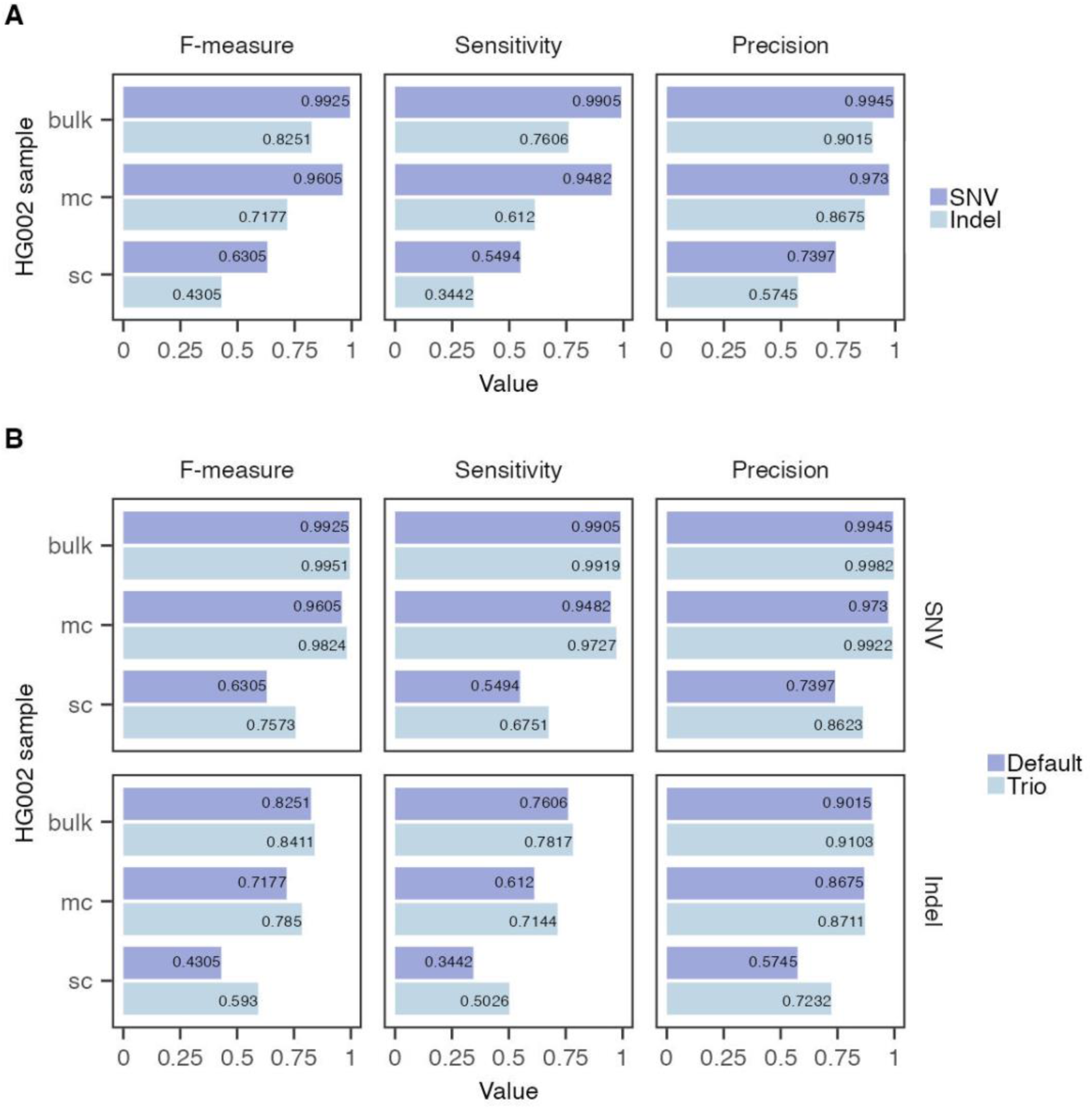
Small variant (SNV and indel) calling performance for lrWGS data from single-cell (sc), multi-cell (mc), and bulk samples of the offspring. (**A**) Variant calling performance in individual samples. (**B**) Trio variant calling yields varying degrees of improvements in variant calling performance for both SNVs and indels. F-measure represents the harmonic mean of precision and sensitivity.

Both single- and multi-cell data contain a high percentage of chimeric reads (55% and 48% for single- and multi-cell data, respectively) **(Supplementary Figure 2A)**. This is expected due to the nature of MDA amplification which is characterized by the formation of chimeric DNA rearrangements. During mapping, each chimeric read was fragmented into multiple smaller segments and mapped to their original positions within the genome.

Among these alignments, one was selected as the representative alignment, while all others were classified as supplementary alignments **(Supplementary Figure 2B)**. By default, supplementary alignments were not utilized for variant calling. We hypothesized that incorporating supplementary alignments might enhance coverage in specific genomic regions and improve SNV calling performance. Hence, we conducted tests that deliberately included supplementary alignments for variant calling. In contrast to expectation, we noted slightly reduced SNV F-measures compared to results obtained without including supplementary alignments **(Supplementary Figure 2C)**. Supplementary alignments were thus not utilized for variant calling throughout this study.

High-quality SNVs and indels are required to enable haplotype phasing. Since both single- and multi-cell data showed lower SNV and indel calling performance compared to bulk data **(Figure 2A)**, we aimed to improve variant calling performance by incorporating parental data (publicly available lrWGS ONT data with ∼24x coverage, see methods) to enable trio information-aware variant calling and reduce Mendelian inheritance violation variants. Trio variant calling was performed using Clair3-Trio (26), resulting in varying degrees of improvements in variant calling performance for the offspring, with the most substantial increase observed for single-cell data **(Figure 2B)**. Parental variant detection performance did not yield obvious benefits from trio variant calling **(Supplementary Figure 3)**.

### Long-read whole-genome sequencing of single cells enables accurate phasing of the human genome

The next step is to achieve accurate phasing. We used the pedigree phasing mode of WhatsHap (28) to achieve high phasing performance by incorporating parental data, combining both read-based and genetic phasing. Meanwhile, we also tested the default phasing mode of WhatsHap (28), which relies solely on read-based phasing. We then compared the trio-based variant calling and phasing results with those obtained using the default settings. We tested phasing with only SNVs and with both SNVs and indels, as these two variant types demonstrated different variant calling performances. When phasing only SNVs, 95-98% of heterozygous SNVs were phased into 2,350, 59,763, and 121,116 blocks, with corresponding block NG50 of 2,964,293, 29,851, and 0 bp, covering 90%, 64%, and 40% of the autosomal regions for bulk, multi-cell, and single-cell data respectively. Adding indels for phasing achieved similar statistics (**Figure 3A-E**). When performing pedigree phasing the outcome was spectacularly improved, especially for multi- and single-cell data (**Figure 3A-E**). The most significant improvement was the generation of chromosome-long haplotypes with one block per autosome, covering 98% of the autosomal region for all data types (**Figure 3C and 3E**). We assessed the accuracy of the phased blocks by comparing them with the phased GIAB benchmark data and obtained switch error rates as an indication of phasing accuracy. A lower switch error rate indicates higher accuracy. We observed higher switch error rates in the phased blocks when phasing both SNVs and indels compared to phasing only SNVs, likely due to lower indel calling performance. Compared to default phasing, pedigree phasing resulted in decreased accuracy for bulk data but improved accuracy for multi- and single-cell data. For bulk data, the switch error rates increased from 0.09% to 0.11% when phasing only SNVs and from 0.22% to 0.49% when phasing both SNVs and indels. In contrast, for multi-cell data, the switch error rates decreased from 0.40% to 0.11% when phasing only SNVs and from 0.54% to 0.44% when phasing both SNVs and indels. For single-cell data, the rates decreased from 1.31% to 0.19% when phasing only SNVs and from 1.46% to 0.50% when phasing both SNVs and indels (**Figure 3F**).

**Figure 3.**
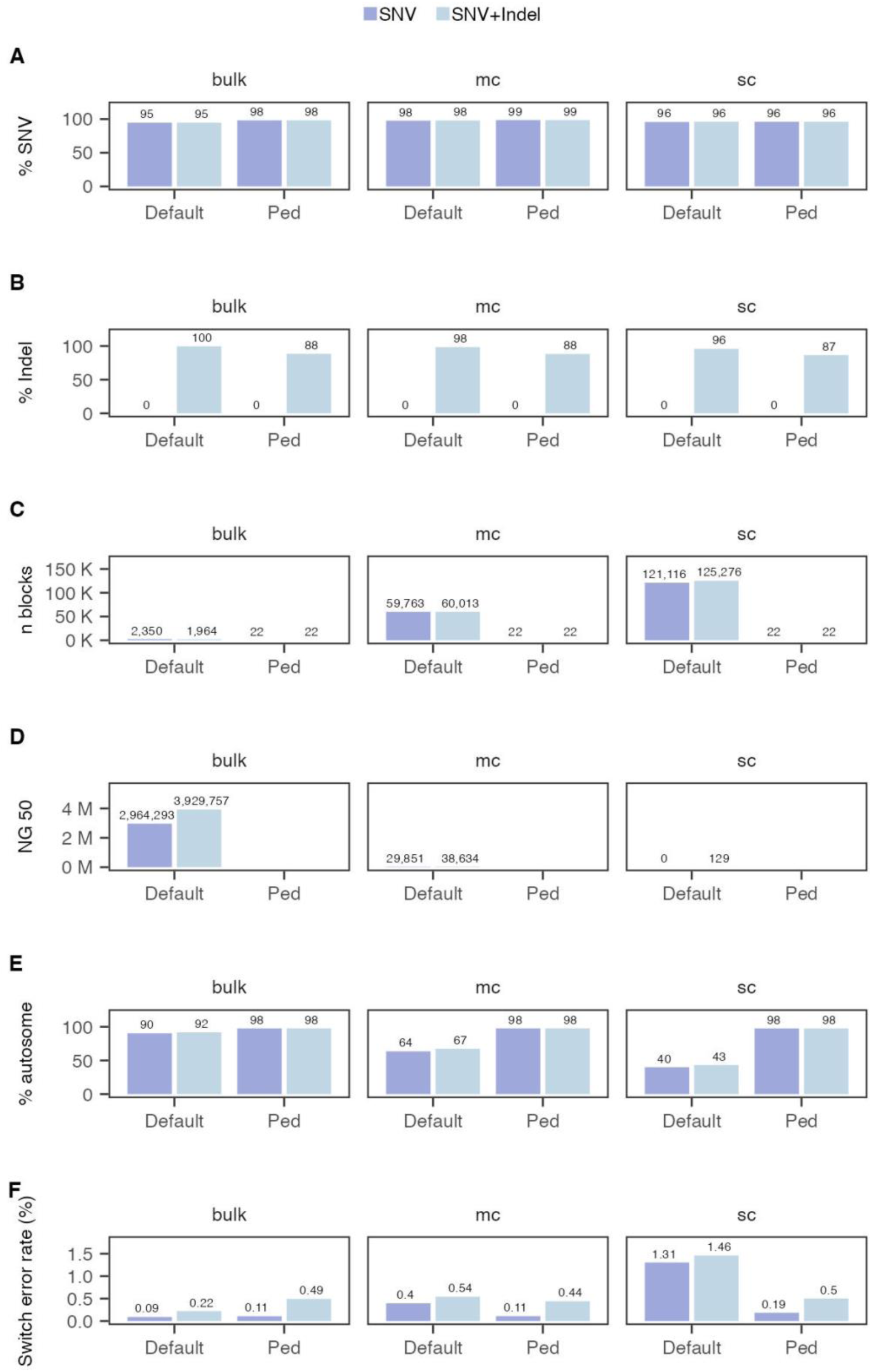
Phasing performance is superior with the pedigree phasing mode (ped) compared to the default phasing mode (default) for single-cell (sc) and multi-cell (mc) data of the offspring. **(A)** Percentage of phased heterozygous SNVs. (**B)** Percentage of phased heterozygous indels. (**C)** Total number of phased blocks. (**D)** NG50 of phased blocks. (**E)** Percentage of autosomal region covered by phased blocks. (**F)** Switch error rate. Shown are statistics for autosomes. For NG50, values are calculated per chromosome and averaged across autosomes. NG50 values are not applicable for pedigree phasing results because each chromosome has only one phased block. For each phasing mode, purple bars represent results when phasing only SNVs, while blue bars represent results when phasing both SNVs and indels.

### Performance comparison of direct variant detection and haplotype-based variant inference

Given the promising quality of trio-based variant calling and phasing from single- and multi-cell data, we hypothesized that the resulting haplotypes could be used to infer parental haplotypes inherited by the offspring, enabling genotype extraction based on these inferred parental haplotypes. To infer the transmitted parental haplotypes, we compared parental haplotypes from default phasing mode with haplotypes of the offspring from pedigree phasing mode. Haplotype comparison was conducted within 1Mb segments across the genome (methods; **Figure 1**). Compared to the transmitted parental haplotypes inferred from the offspring’s bulk data phasing results, those inferred from the offspring’s multi- and single-cell data phasing results demonstrated consistencies of 95% and 82% when phasing only SNVs, and 93% and 82% when phasing both SNVs and indels, respectively **(Supplementary Figure 4)**.

We then assessed the specific added value of using long reads for phasing. With short-read or SNP arrays data, genetic phasing is typically applied within a pedigree. This method can phase variant loci that are heterozygous in one parent and homozygous in the other, as well as loci exhibiting different homozygous genotypes in the parents. However, loci that are heterozygous across all trio members remain unresolved. In contrast, long-read data have the potential to phase these loci. We evaluated the proportions of these loci among all loci used for haplotype comparison (**Supplementary Figure 5**). The results showed that 15-19% of informative SNVs and 14-17% of informative indels were heterozygous across trio members and could only be phased with long reads (**Supplementary Figure 5**). This indicates that lrWGS data enable more loci to be phased and utilized for subsequent analyses.

Next, we compared the performance of two variant detection approaches: direct variant detection and haplotype-based analysis. In direct variant detection, variants are identified from default variant calling results, whereas in haplotype-based analysis, variants are inferred from the transmitted parental haplotypes **(Figure 1)**. Using GIAB benchmark data, we selected loci within high-confidence protein-coding gene regions where one parent is homozygous reference (0/0) and the other is heterozygous (0/1). The offspring can be either heterozygous (0/1) or homozygous reference (0/0), allowing us to evaluate the performance of both approaches in detecting transmitted coding variants. In total, we selected 812,973 SNV loci, with 407,232 heterozygous and 405,741 homozygous reference in the offspring, and 102,465 indel loci, with 50,650 heterozygous and 51,815 homozygous reference in the offspring. Using direct variant detection, we correctly identified 94% of heterozygous SNVs in multi-cell data and 43% in single-cell data, with no false positives for homozygous reference loci (**Figure 4A**). Performance for indel loci was lower, with 68% of heterozygous indels detected in multi-cell data and 26% in single-cell data, along with 1% false positives for homozygous reference loci (**Figure 4B**). In contrast, haplotype-based analysis demonstrated superior performance over direct variant detection, accurately inferring 98% of heterozygous SNVs in multi-cell data and 92% in single-cell data, with 2-3% false positives for homozygous reference loci (**Figure 4A**). The performance for indel loci was still lower, identifying 78% of heterozygous indels in multi-cell data and 74% in single-cell data, with 4-5% false positives for homozygous reference loci (**Figure 4B**). Multi-cell data demonstrated direct variant detection performance comparable to bulk data and also benefited from haplotype-based analysis, identifying more heterozygous loci despite a small rise in false positives for homozygous reference loci (**Figure 4**).

**Figure 4.**
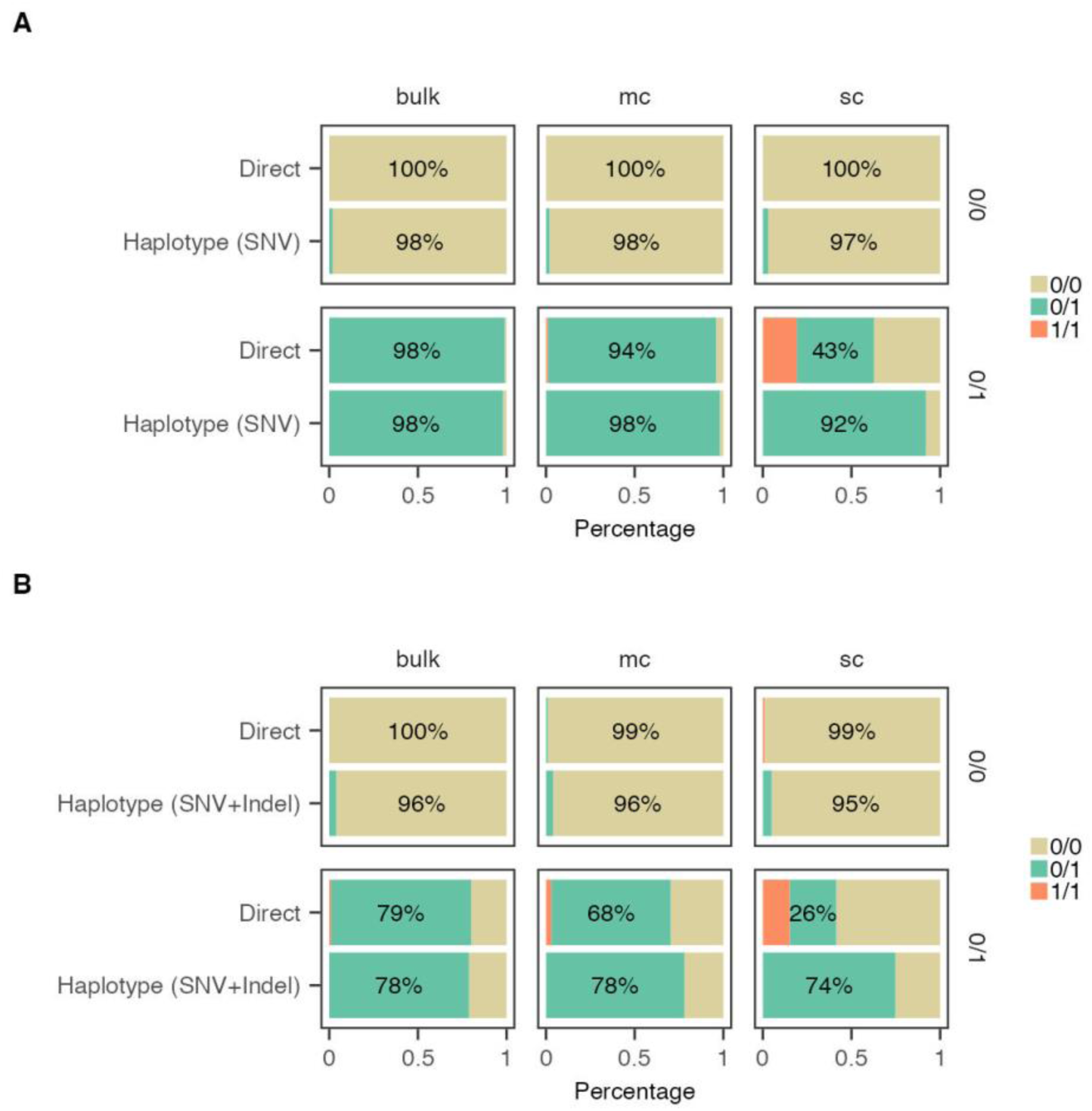
Performance comparison of direct variant detection and haplotype-based variant inference on selected (A) SNV and (B) indel loci using single-cell (sc), multi-cell (mc), and bulk data of the offspring. Direct: direct variant detection. Haplotype (SNV): haplotype-based analysis with only SNVs phased in the haplotypes. Haplotype (SNV + Indel): haplotype-based analysis with both SNVs and indels phased in the haplotypes. The true genotypes from GIAB benchmark data are indicated on the right. The proportions of inferred genotypes are color-coded. 0/0 for homozygous reference, 0/1 for heterozygous, and 1/1 for homozygous alternate.

### Exploration of de novo mutation screening

An essential aspect of single-cell genomics is exploring new mutations that arise in individual cells during processes such as cell division, cancer genesis and tissue differentiation. Understanding DNM rates at the single-cell level will deepen our knowledge of the heterogeneity of both cancerous and normal cells, as well as the mechanisms driving cancer progression. Additionally, a large proportion of common and rare genetic disorders are a consequence of DNMs (Veltman and Brunner, 2012). Therefore, we assessed the efficacy of lrWGS data for genome-wide DNM screening, focusing on SNVs. We identified 452 heterozygous SNVs in the offspring as “true DNMs” within high-confidence protein-coding regions of the GIAB benchmark data, with both parents exhibiting homozygous reference genotypes at the corresponding loci. This is significantly higher than expected, since the number of new point mutations present in human offspring is on average 60 (30–90 depending on parental age at conception), with around one DNM per exome (31–34). Hence, most are likely false positives or mosaic variants introduced by cell culture. Of the 452 DNMs detected in the benchmark data, 407 (90%) were identified using multi-cell data, while 167 (37%) were detected with single-cell data from the offspring. However, the false positive rates were 87,434 out of 87,841 (99.5%) for multi-cell data and 321,201 out of 321,368 (99.9%) for single-cell data (**Supplementary Table 4**). In summary, the abundance of false positives complicates DNM screening when using single-cell or multi-cell lrWGS data of the offspring for DNM screening.

### Direct variant detection, haplotype-based variant inference, and haplotype-aware aneuploidy profiling in human embryos using lrWGS data

Since the proof-of-concept study with GIAB cell lines demonstrated promising performance, we proceeded to test lrWGS-based PGT on TE biopsies from five human embryos derived from two different couples. Each TE biopsy contains 5-10 cells, corresponding to the multi-cell sample in above benchmark study. The DNA was previously analyzed using clinically accredited SNP array-based comprehensive PGT, and the results were used as a reference (**Supplementary Table 5 and Supplementary Figure 6**). For family ONT1 the father carried a pathogenic SNV (c.1384C>T) in the *MSH2* gene causing Lynch syndrome, an autosomal dominant cancer predisposition syndrome. For family ONT2, the mother carried an indel (c.2955delG), and the father carried a pathogenic SNV (c.1133-708A>G) in the *LAMB3* gene, responsible for autosomal recessive junctional epidermolysis bullosa. Two embryos from family ONT1 (ONT1-E02, ONT1-E03) and three from family ONT2 (ONT2-E04, ONT2-E06, ONT2-E20) were tested using lrWGS-based PGT. We obtained 21-31x coverage following lrWGS of the parents and embryos, covering 93-95% of the human genome (**Supplementary Table 6 and 7**). With direct mutation detection, the carrier status of the variant alleles was accurately determined, except for one indel that was missed in ONT2-E20. However, we identified 3 out of 5 reads supporting the presence of this deletion (**Supplementary Table 8**). Haplotype-based analysis yielded 100% concordance with SNP array-based PGT-M results **(Figure 5)**. SNVs were phased to identify the risk parental haplotypes carrying the disease alleles and to infer the parental haplotypes inherited by the embryo, from which the embryo’s carrier status can be determined (**Figure 5**). For the indel in the mother of family ONT2, the risk haplotype carrying the indel was identified by phasing both the indel and the SNVs. The indel locus was then incorporated into the SNV phasing results, with each allele assigned to the corresponding maternal haplotype (**Figure 5B**).

**Figure 5.**
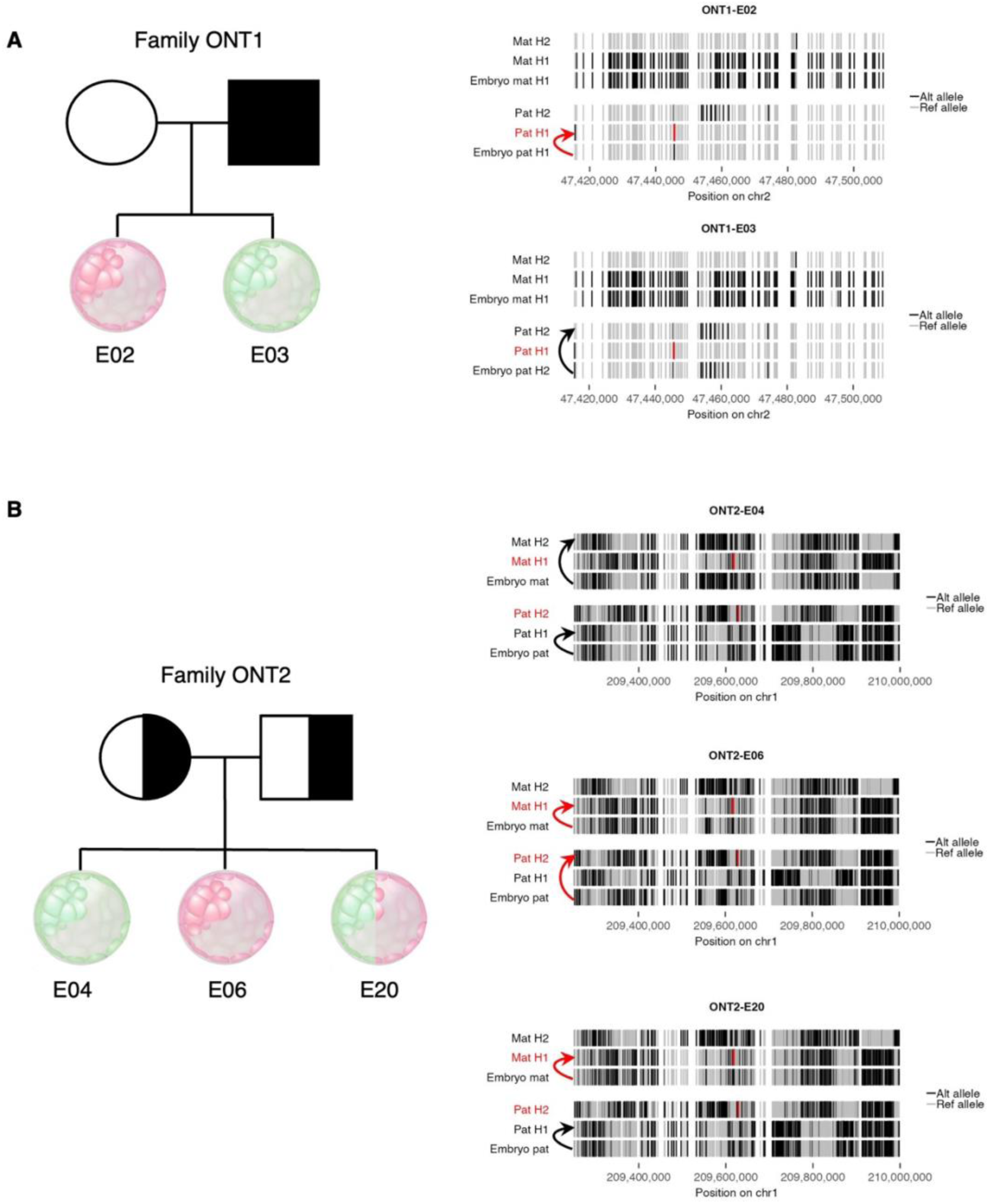
Visualization of haplotype-based PGT-M in human embryos. Pedigree plots for the two PGT families are shown on the left. Embryos are color-coded based on their carrier status as determined by haplotype-based PGT-M: green indicates that the disease allele was not inherited, red indicates that the dominant mutation was inherited or both recessive mutations were inherited, and green/red mosaic indicates the inheritance of a single recessive mutation. SNV haplotyping results for the chromosomal region linked to the disease loci are shown on the right. In the maternal (Mat H1, Mat H2) and paternal (Pat H1, Pat H2) haplotypes, each vertical line represents an allele from an informative locus: grey indicates the reference allele, black indicates the alternate allele, and disease alleles are shown in red. Parental haplotypes that carry disease alleles are labeled in red text. Arrows link the embryonic haplotypes to their corresponding maternal and paternal haplotypes, with red arrows used when the corresponding parental haplotypes carry the disease alleles. For the indel in the mother of family ONT2, both SNVs and indels were first phased to obtain phasing information for the indel locus. This locus was then added to the SNV phasing results, with each allele assigned to its corresponding haplotype. (**A**) In Family ONT1, ONT1-E02 inherited the paternal haplotype carrying the disease allele, while ONT1-E03 inherited the normal paternal haplotype. (**B**) In Family ONT2, ONT2-E04 inherited normal haplotypes from both parents, ONT2-E06 inherited pathogenic allele-carrying haplotypes from both parents, and ONT2-E20 inherited the normal haplotype from the father and the pathogenic allele-carrying haplotype from the mother.

Since SNP array-based comprehensive PGT allows for the concurrent analysis of aneuploidy (PGT-A) and enables the identification of mosaic aneuploidy and the origin of aneuploidy, we explored whether these can also be detected by lrWGS data. We identified all aneuploidies and their mechanistic origins using lrWGS data, achieving 100% consistency with SNP array-based PGT (**Figure 6 and Supplementary Figure 6 and 7**). Specifically, in ONT1-E02 we observed complex segmental aberration on chromosome 2, with mosaic duplication of short arm and mosaic deletion of the long arm. In addition, the same embryo exhibited complex segmental aberrations on chromosome 14 (**Supplementary Figure 7**). Additionally, we detected trisomy on chromosomes 9 and 16 in ONT2-E20 **(Figure 6A)**, with parental allele fractions indicating the presence of an extra maternal chromosome **(Figure 6B)**. To determine the mitotic or meiotic origin of the extra maternal chromosomes, we analyzed loci with homozygous genotypes in the father and heterozygous genotypes in the mother and the embryo. For each locus, we calculated the unique allele fraction (UAF) in the embryo, with the unique allele being the distinct allele among the four parental alleles. The distribution of the unique allele fraction across the chromosome helps identify the mechanistic origin of the trisomy **(Figure 6C)**. Both chromosomes 9 and 16 were inferred to originate from maternal meiotic I nondisjunction **(Figure 6D)**, consistent with SNP array results **(Supplementary Figure 6E)**.

**Figure 6.**
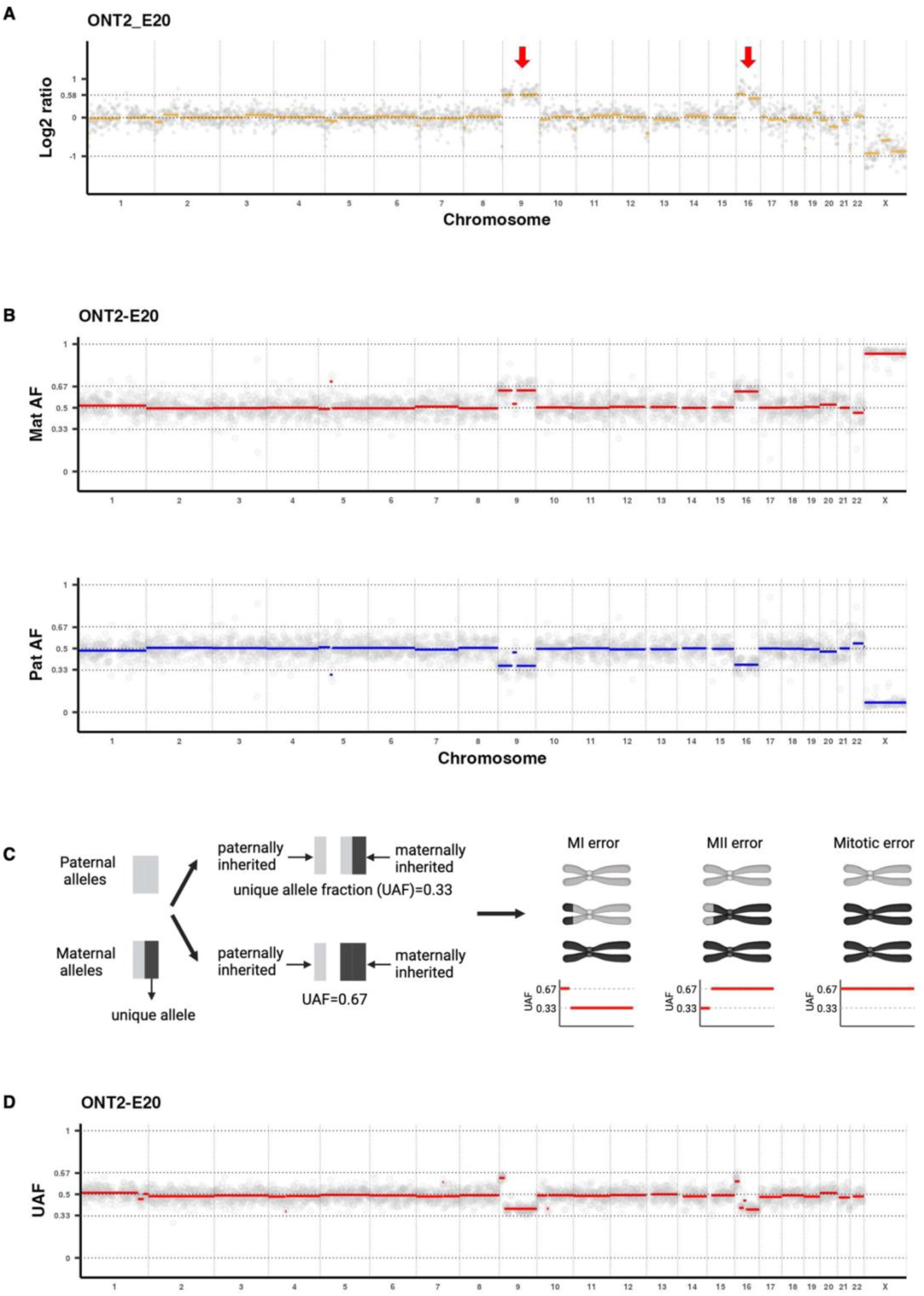
PGT-A analysis using lrWGS data identified aneuploidies and their mechanistic origins in ONT2-E20. (**A)** The copy number plot shows trisomy on chromosomes 9 and 16. (**B**) Paternal allele fractions (Pat AF) and maternal allele fractions (Mat AF) were assessed across the genome for loci homozygous in one parent and heterozygous in the other, reflecting parental genome contributions across the embryonic genome. In the trisomic regions, Mat AF = 0.67 and Pat AF = 0.33, indicating one additional chromosome from the maternal side for chromosomes 9 and 16. (**C**) Schematic illustrating the inference of mechanistic origins for maternally derived trisomies. Loci with homozygous genotypes in the father and heterozygous genotypes in both the mother and the embryo are selected. One locus is shown, with grey and black bars representing different alleles; in this case, the black bar represents the unique allele among the four parental alleles. The unique allele fraction (UAF)—the fraction of this unique allele in the embryo for this locus—is inferred from the allele frequency (AF) value in variant calling results, which is expected to be 0.33 if two different maternal alleles were inherited and 0.67 if two identical maternal alleles were inherited. The pattern of UAF values across the trisomic chromosome indicates the mechanistic origin of the trisomy. (**D**) UAF values across the genome. For trisomic chromosomes 9 and 16, the UAF is 0.67 at the beginning and decreases to 0.33 across the remaining chromosomal regions. This pattern indicates that the origin of these two trisomies is maternal meiotic I nondisjunction.

In summary, we confirmed in human embryos that lrWGS can be used for concurrent PGT-M and PGT-A analysis. For PGT-M, while direct variant detection missed an inherited indel in one of the embryos, inclusion of haplotype-based analysis mitigated this drawback, resulting in 100% concordance with SNP array-based PGT results. For PGT-A, lrWGS data enables not only the detection of aneuploidies but also their parental and mechanistic origins.

## Discussion

Here, we explored the characteristics of lrWGS data from single cells and evaluated its performance for variant calling and reference-free haplotyping. Our benchmark analysis revealed that with lrWGS data from TE biopsies (10 cells), direct mutation analysis has a 94% probability of identifying an inherited SNV. For haplotype-based analysis, the probabilities are 98% for TE biopsies (10 cells) and 92% for single blastomere biopsies (1 cell). Considering that TE biopsy is becoming the golden standard, lrWGS-based PGT can thus enable direct variant detection coupled with haplotype-based analysis for increased diagnostic accuracy. Using human embryos, we validated the high performance of lrWGS-based PGT-M and concurrent PGT-A, with the mechanistic origins of aneuploidies correctly identified.

Correctly matching the haplotypes of the offspring to those of their parents is essential for effective haplotype linkage analysis. Key factors contributing to successful matching include: 1) Attainment of high-quality variant calling and phasing outcomes for the parents, which played a pivotal role in identifying high-risk and low-risk haplotypes. 2) Trio variant calling for the offspring predicted significantly fewer Mendelian inheritance violation loci. 3) Pedigree-based phasing for the offspring generated chromosome-spanning haplotypes with increased phasing accuracy by combining read based phasing and genetic phasing. 4) The increased density of informative loci in our study, primarily due to loci that could only be phased using long reads, enabled successful haplotype matching over relatively short genome regions. 5) Exclusion of loci violating Mendelian rules from haplotype comparison. 6) Flexibility in the haplotype matching process, allowing for discrepancies between the offspring’s haplotype and the inferred parental haplotype.

The main advantage of using parental lrWGS data for phasing is that it allows for direct phasing of the parents, thereby enabling genetic phasing of single cells from the offspring without requiring additional family members. However, genetic phasing within a trio cannot resolve loci where all trio members are heterozygous (15). By incorporating lrWGS data from single cells of the offspring, we found that 14-19% of informative loci fall into this category. These additional informative loci increase the probability that loci affected by ADO in single-cell data can be correctly inferred from inherited parental haplotypes by enhancing the density of informative loci available for haplotype matching. The broad genome coverage of lrWGS data from single cells also enables direct variant detection. These advantages make lrWGS-based PGT a superior option compared to current PGT methods. Firstly, there is no need to obtain DNA from close relatives, which often causes delays, increases costs, and may not always be available. Secondly, lrWGS-based PGT enables the direct phasing of DNMs present in prospective parents , eliminating the need for additional steps such as analyzing single sperm, polar bodies (35–37) or affected sibling embryos(38). Thirdly, it allows for the direct detection of pathogenic variants in embryos, complementing haplotype-based analysis by resolving uncertain or inconclusive findings and addressing cases where the embryo is at risk of carrying a pathogenic mutation due to parental germline mosaicism. Fourthly, using a higher density of informative loci for analysis enhances the identification of inherited parental haplotypes within relatively short regions. Additionally, lrWGS data from single cells allows for the detection of both the parental origin and the mitotic or meiotic nature of chromosomal anomalies, providing valuable insights into the etiology of aneuploidies. Such information is crucial in clinical practice, as aneuploidies resulting from meiotic chromosome segregation errors rarely survive to term and often lead to adverse pregnancy outcomes; thus, selecting against these embryos could improve IVF success rates (39). The potential applications of lrWGS-based single-cell haplotyping and aneuploidy profiling go beyond human PGT and can be adapted for other species, such as equine and bovine, to improve reproductive outcomes. It also holds promise for cell-based noninvasive prenatal diagnosis by analyzing single fetal cells present in maternal blood (40).

DNMs arise during various biological and pathological processes, such as cell division and cancer development. Additionally, DNMs are a major cause of rare human disorders. Genome-wide DNM screening would be valuable for identifying these mutations. Using multi- and single-cell lrWGS data of the offspring, we identified 90% and 37% of DNMs in benchmark data, respectively. However, true DNMs represented only 0.5% and 0.1% of all identified DNM candidates, highlighting the abundance of false positives. These results demonstrate that whole-genome DNM screening with ONT lrWGS data remains a challenging task at present However, with increasing sequencing accuracy and methodological improvements, this is likely to be possible in the future. A previous pilot study attempted to use variant annotation databases and functional prediction algorithms to identify real pathogenic DNMs among numerous DNM candidates (41). Such strategies and additional quality metrics could be integrated into DNM screening to enhance detection accuracy. It is worth noting that we used cell line samples for DNM detection. Since DNMs arise during each cell division and increase with each passage of culturing (Londin et al., 2011), the culture process may have influenced the observed high incidence of DNMs.

In this study, we achieved high-quality phasing results with lrWGS data from the bulk sample of the offspring. Compared to a previous study that phased the autosomes of the same individual into 19,215 blocks with a switch error rate of 0.37% for SNVs and indels using 28x PacBio HiFi reads (5), we obtained 1,964 phased blocks with a lower switch error rate of 0.22% using ∼24x ONT reads. This improvement is notable despite the lower variant calling performance of ONT reads compared to HiFi reads (F-measures: 0.9991 for SNVs and 0.9598 for indels with HiFi reads; 0.9925 for SNVs and 0.8251 for indels with ONT reads). The improvement in phasing performance could be attributed to advancements in the reference genome, updates in bioinformatics software, and enhancements in the GIAB benchmark data, as newer versions were utilized for this study.

A constant recombination rate of 1.26 cM/Mb was used during phasing, which is a suitable assumption for the human genome. However, the inherent errors in lrWGS data, especially those derived from single-cell and multi-cell samples, may hinder the accurate detection of actual recombination sites. Consequently, unidentified recombination events may affect haplotype comparison. Moreover, imperfect phasing results contain switch errors that may influence haplotype matching. To mitigate the impact of recombination events and switch errors on the inference of inherited parental haplotypes, we manually constrained the maximum comparison block length to 1 Mb in this study. Additionally, visual inspection of haplotype blocks can help identify recombination events and switch errors, further reducing the risk of misdiagnosis. With ongoing improvements in sequencing accuracy, read length, and bioinformatics algorithms, this constraint on block length may eventually become unnecessary, making accurate phasing of the entire genome feasible.

This study has limitations and areas for potential improvement. First, we utilized ONT sequencing, a cost-effective long-read sequencing technology that is rapidly advancing in read length and accuracy. These improvements will enhance variation discovery and phasing performance, making it essential to conduct updated benchmark studies regularly. Second, more data from clinical applications are needed to further validate the practical utility of lrWGS-based haplotyping and aneuploidy profiling in single cells. As lrWGS becomes more widely adopted in clinical settings, performance evaluations from large-scale clinical PGT cycles could provide additional insights into the efficiency of lrWGS-based PGT in a broader range of clinical scenarios.

To summarize, we developed a bioinformatics pipeline that enables genome-wide concurrent haplotyping and aneuploidy profiling of single cells using lrWGS data, and validated its effectiveness for genome-wide, reference-free comprehensive PGT. Additionally, we evaluated the performance of lrWGS data from single cells for direct variant detection and DNM screening. Beyond PGT for human embryos, our bioinformatics pipeline has potential applications in other areas of single-cell genomics. For instance, it can be adapted for PGT in animal species like bovine and equine to improve reproductive outcomes, and for cell-based noninvasive prenatal testing by analyzing single circulating trophoblast cells in maternal blood (40).

## Data availability

Raw data has been deposited at the European Genome-phenome Archive (EGA), which is hosted by the EBI and the CRG, under accession number EGAD50000000787. It is available to academic users upon request to the Data Access Committee (DAC) of KU Leuven via the corresponding author (JRV). We have provided the bioinformatical scripts via the following link: https://github.com/JorisVermeeschLab/ONT_PGT.git

## Supplementary data

Supplementary data are available at NAR online.

## Acknowledgements

The authors would like to thank the couples who participated in the study.

