## Supplementary Figure 1-7 for "Long-read whole-genome sequencing-based concurrent haplotyping and aneuploidy profiling of single cells"

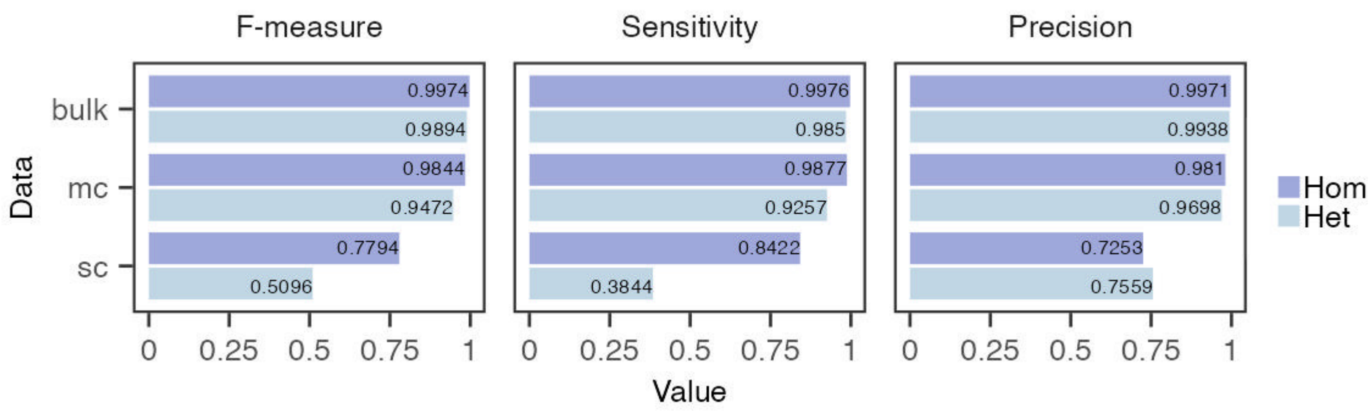


**Supplementary Figure 1.** Heterozygous and homozygous SNV calling performance for lrWGS data from single-cell (sc), multi-cell (mc), and bulk samples of the offspring.

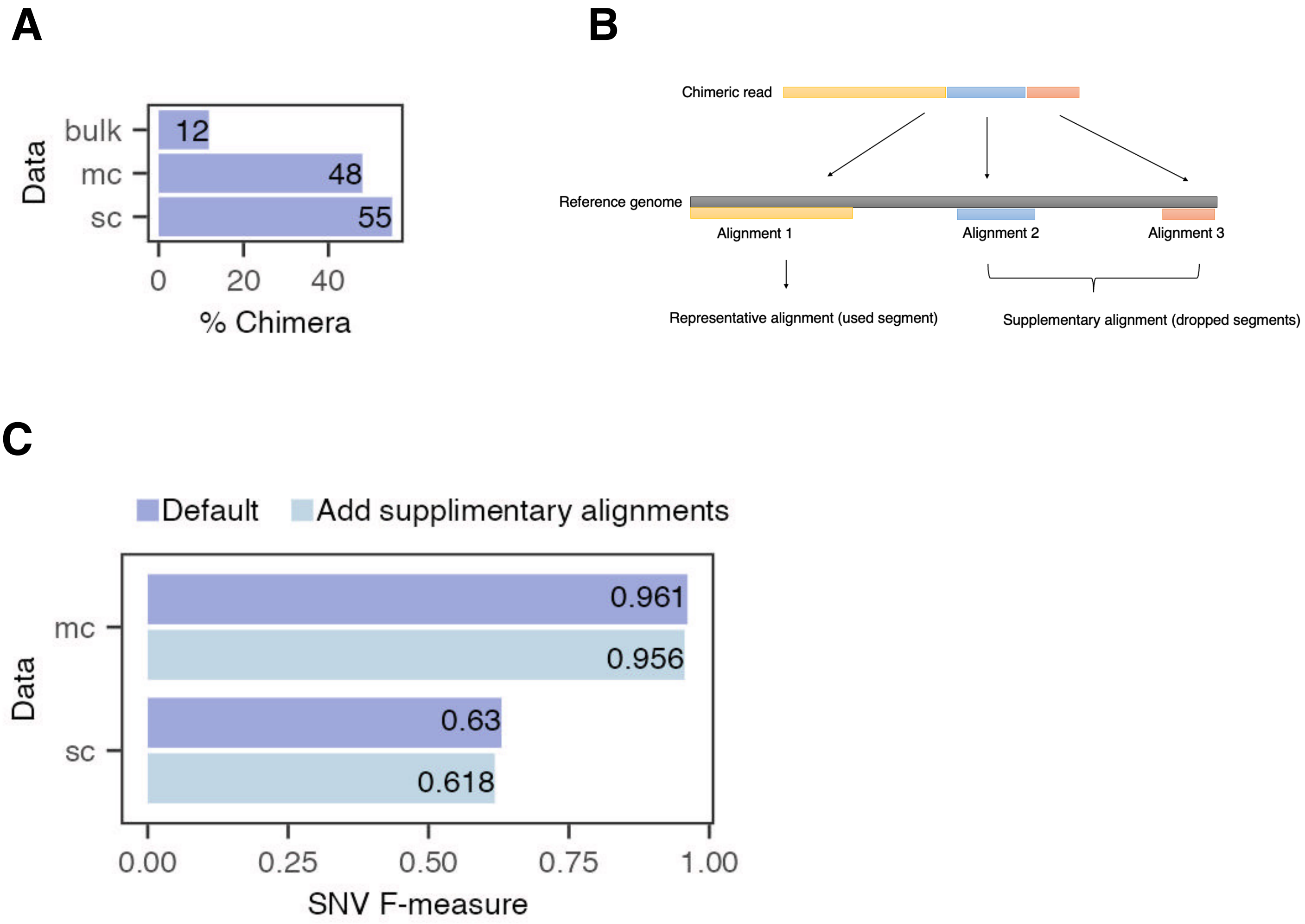


**Supplementary Figure 2.** Supplementary alignments did not improve SNV calling performance. **(A)** Percentage of chimeric reads in bulk, multi-cell (mc) and single-cell (sc) lrWGS data of the offspring. **(B)** Schematic demonstration of chimeric reads and supplementary alignments. **(C)** Comparison of default SNV calling F-measures with those obtained after including supplementary alignments for variant calling.

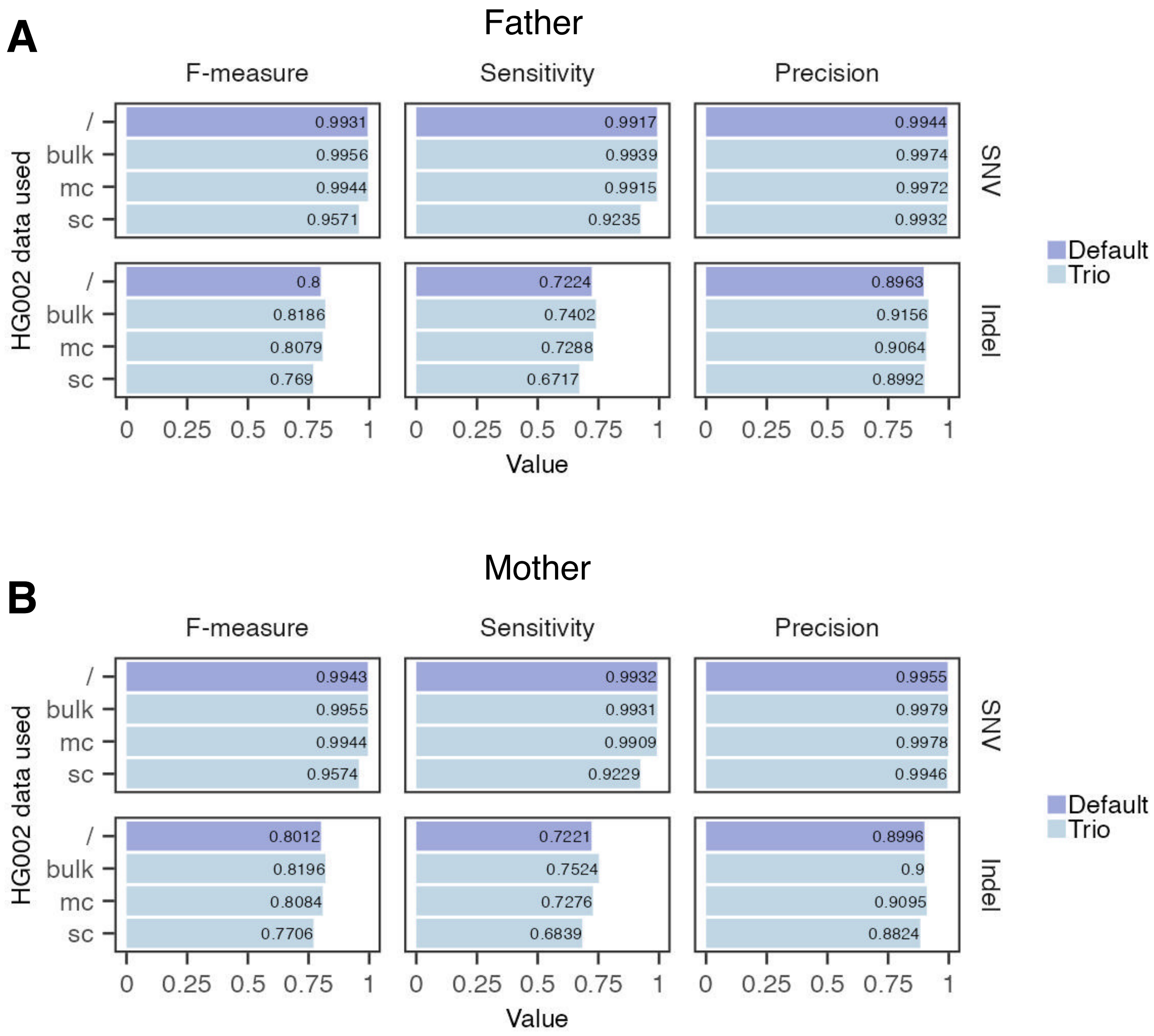


**Supplementary Figure 3.** Small variant (SNV and indel) calling performance obtained form default and trio variant calling for **(A)** the father and (**B**) the mother. On the y-axis, "bulk," "mc," and "sc" indicate bulk, multi-cell (mc), or single-cell (sc) data of the offspring were used for trio variant calling together with parental data. The "/" denotes only paternal or maternal data were utilized for variant calling.

**A**

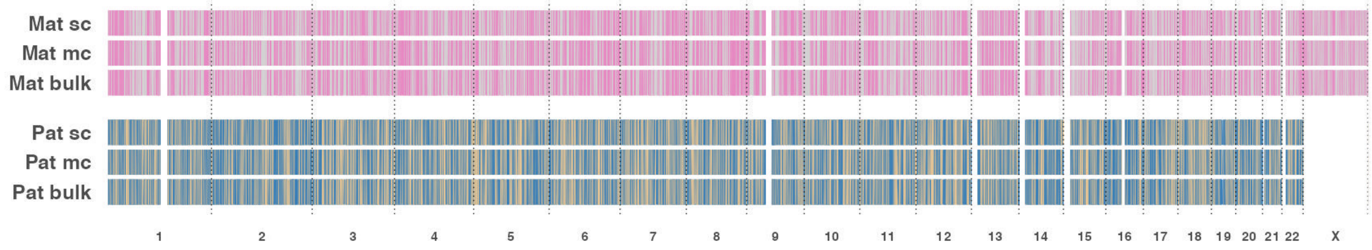


**B**

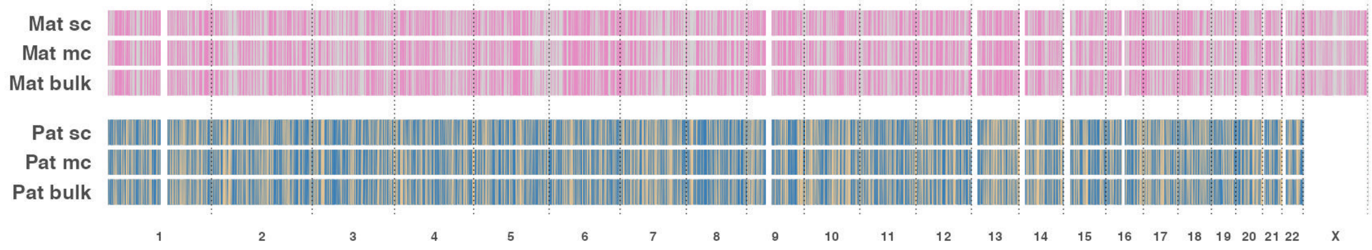


**Supplementary Figure 4.** Consistency of transmitted parental haplotypes inferred from the offspring's bulk, multi-cell (mc) and single-cell (sc) data phasing results when phasing **(A)** only SNVs **(B)** SNVs and indels. For each panel, vertical colored lines indicate inferred paternal (pat) or maternal (mat) haplotypes for specific regions. Different parental haplotypes are distinguished by different colors.

**A**

| Trio genotype combinations for informative loci | | | Phasing method |
|---|---|---|---|
| Father | Mother | Offspring | |
| 0\|1 or 1\|0 | 0\|1 or 1\|0 | 0\|1 or 1\|0 | Read only |
| 0/0 or 1/1 | 0\|1 or 1\|0 | 0\|1 or 1\|0 | Genetic or read |
| 0\|1 or 1\|0 | 0/0 or 1/1 | 0\|1 or 1\|0 | Genetic or read |
| 0/0 | 1/1 | 0\|1 or 1\|0 | Genetic or read |
| 1/1 | 0/0 | 0\|1 or 1\|0 | Genetic or read |

**B**

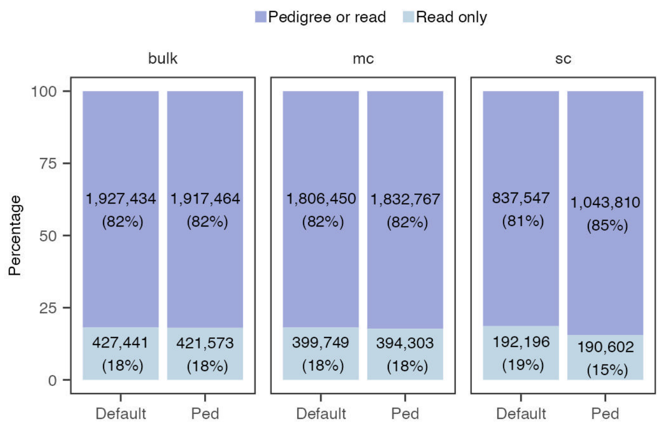


**C** **D**

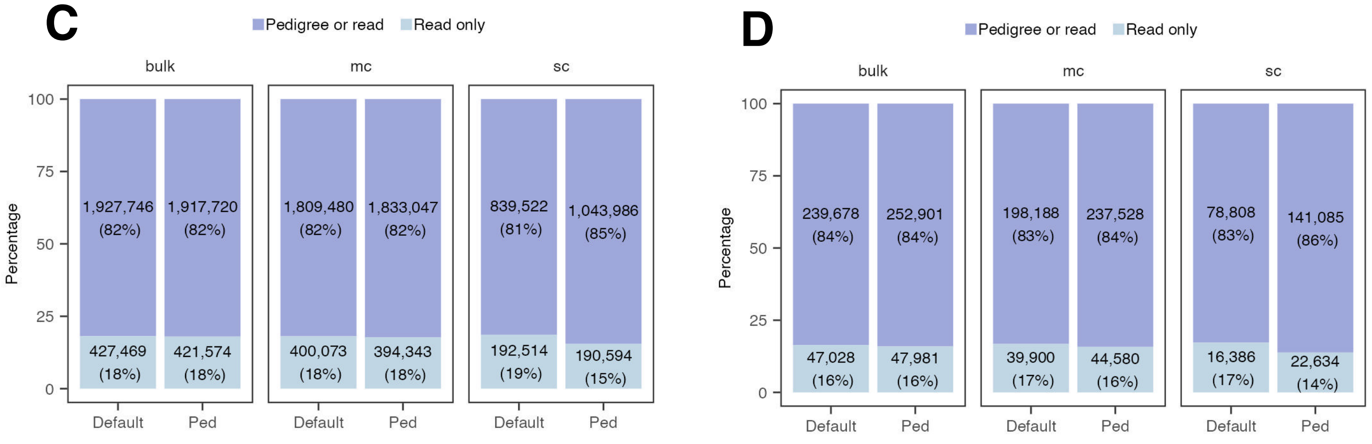


**Supplementary Figure 5.** Proportion of informative loci that can only be phased by long reads in pedigree (ped) or default phasing (default) results of bulk, multi-cell (mc) and single-cell (sc) data. **(A)** Trio genotype combinations for loci retained as informative loci for haplotype comparison. The "Phasing method" column indicates method(s) that could be used to phase variants loci of the offspring. Read only: variant loci could only be phased through read-based phasing. Genetic or read: variant loci could be phased by either genetic phasing or read-based phasing. **(B-D)** Informative **(B)** SNVs from SNV phasing results, **(C)** SNVs from SNV and indel phasing results, and **(D)** indels from SNV and indel phasing results retained for haplotype comparison, categorized by phasing method(s) that could be used to phase the variant loci.

**Supplementary Figure 6.** SNP array-based comprehensive PGT results for **(A)** ONT1-E02, **(B)** ONT1-E03, **(C)** ONT2-E04, **(D)** ONT2-E06 and **(E)** ONT2-E20. For each embryo, the first panel displays genome-wide maternal and paternal haplarithm plots (Mat-BAF and Pat-BAF tracks) along with a copy number plot (logR track) for chromosomes 1 to X. For family ONT1, haplotyping was performed with paternal grandparents, and thus, only the paternal haplarithm is shown. For family ONT2, haplotyping was performed with a sibling, allowing both maternal and paternal haplarithms to be displayed. Aneuploidies are identified via the logR track, with chromosomal gains marked by red rectangles and chromosomal losses by green rectangles. Combining this information with the paternal and maternal haplarithms helps determine both the parental and mechanistic origins of the aneuploidies. For disomic chromosomes, a distance of 0.5 between the red and blue lines for paternal and maternal haplarithms is expected. Pure trisomy is indicated by a distance of 0.67 in one parent's haplarithm and 0.33 in the other parent's haplarithm. The parent with a distance of 0.33 contributed two chromosomes: when the red and blue lines are at 0 and 0.33 or at 0.67 and 1, this indicates two identical haplotypes from that parent, whereas a distance of 0.33 and 0.67 signifies two different haplotypes from that parent. This information helps identify the parental and mechanistic origin of the trisomy. For mosaic aneuploidies, an increased distance (>0.5) between red and blue lines on one parent's haplarithm suggests a lower chromosomal contribution from that parent, while a decreased distance (<0.5) on the other parent's haplarithm indicates a higher chromosomal contribution from that parent. A comprehensive visual guide for interpreting haplarithm plots is detailed in [1]. The following panel provides a zoomed-in view of the chromosome containing the variant loci of interest. Haplotype blocks are shown, with dark and light blue representing paternal haplotypes and dark and light red representing maternal haplotypes. Dark blue/red indicates the paternal/maternal haplotype carrying the variant allele. Variant loci are marked by vertical orange dashed lines, and the inherited parental haplotypes for these loci are those crossed by the lines, allowing for the inference of the embryo's carrier status. At the bottom, conclusions for PGT-M and PGT-A analyses are provided.


1. Zamani Esteki, M. *et al.* Concurrent Whole-Genome Haplotyping and Copy-Number Profiling of Single Cells. *Am J Hum Genet* **96**, 894 (2015).

**A** ONT1-E02

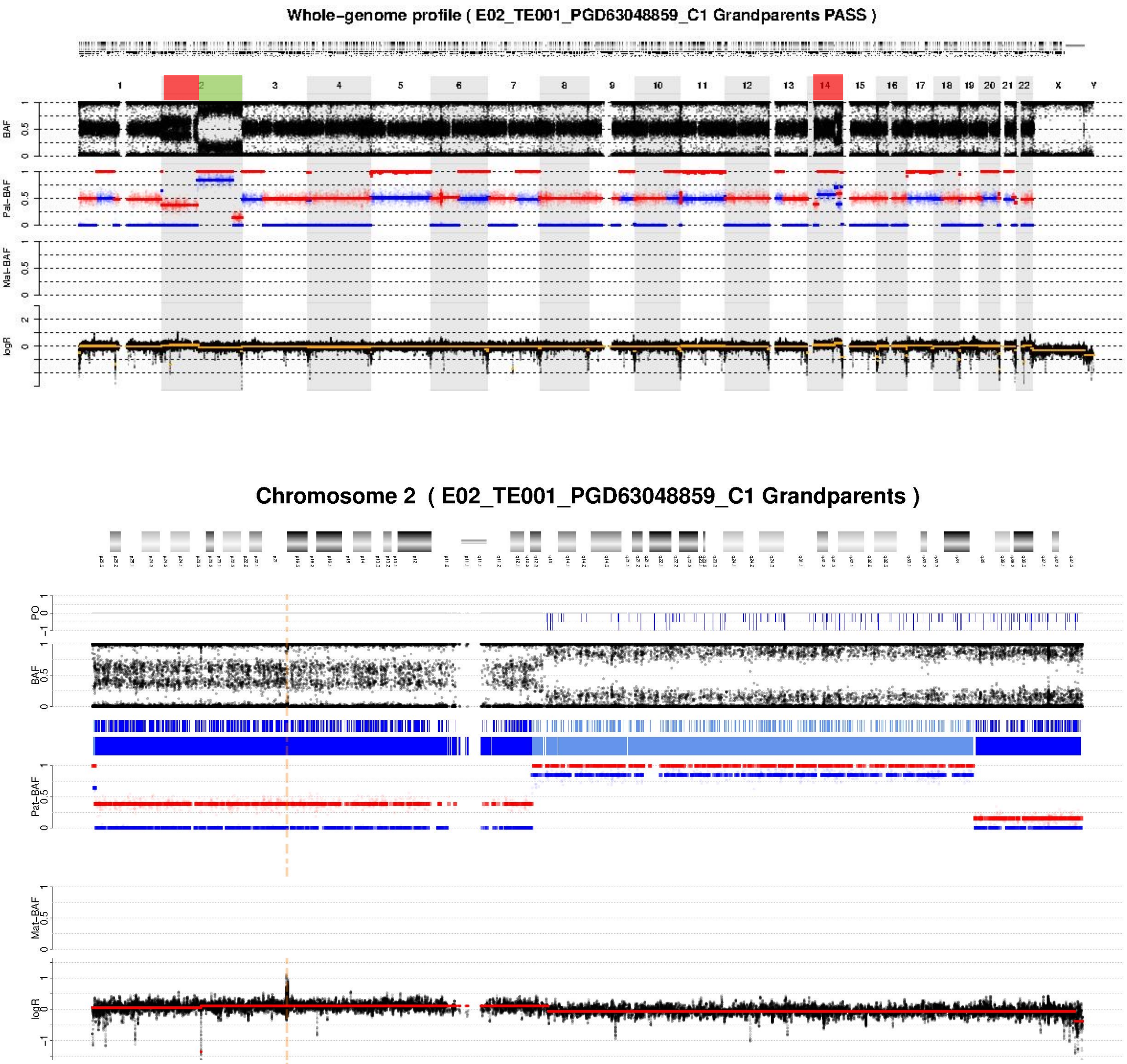


PGT-M:
Paternal SNV in *MSH2*: inherited

PGT-A:
Chromosome 2 mosaic duplication and deletion;
Chromosome 14 mosaic duplication

**B** ONT1-E03

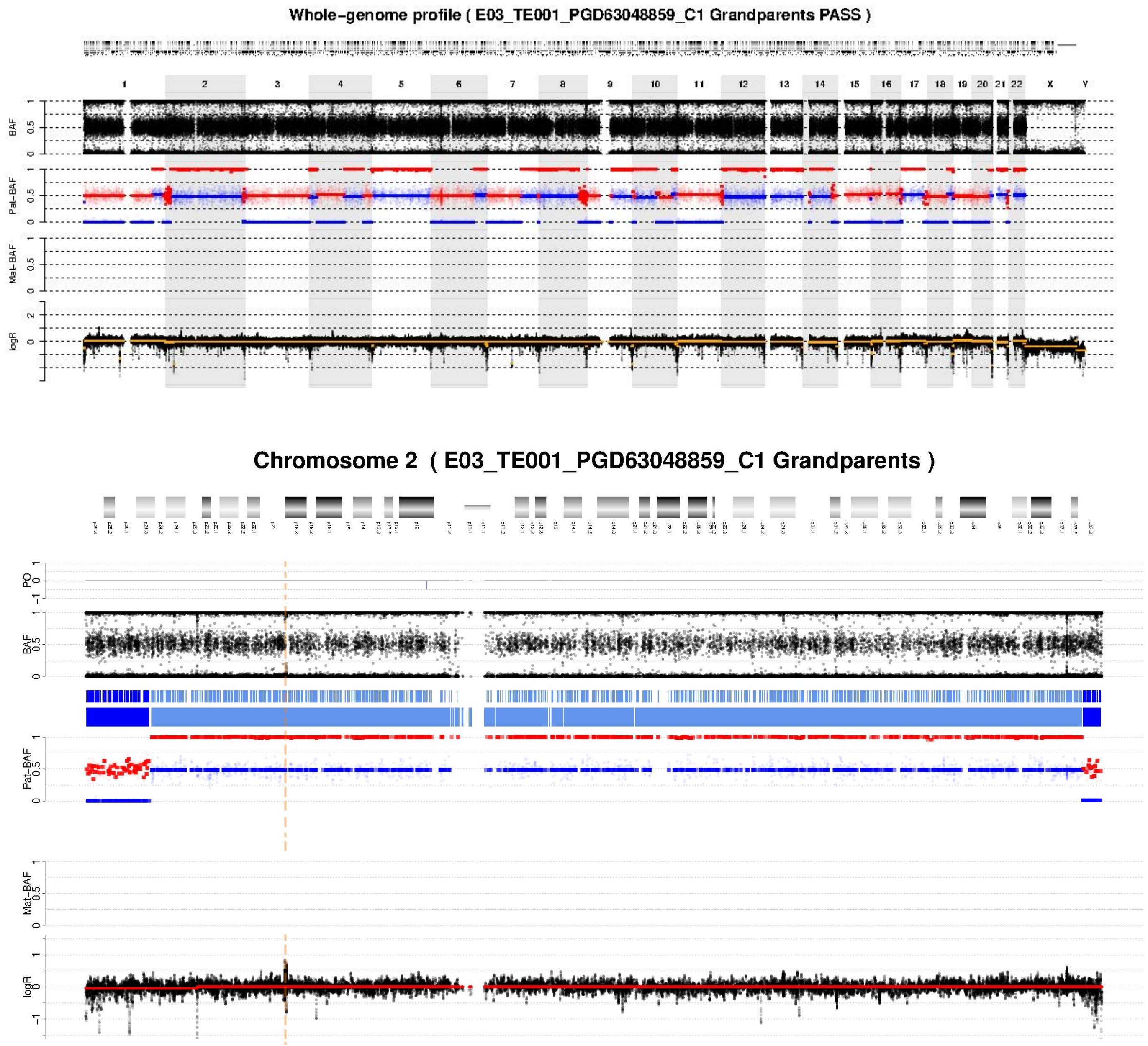


PGT-M:
Paternal SNV in *MSH2*: not inherited

PGT-A:
Euploid

**C** ONT2-E04

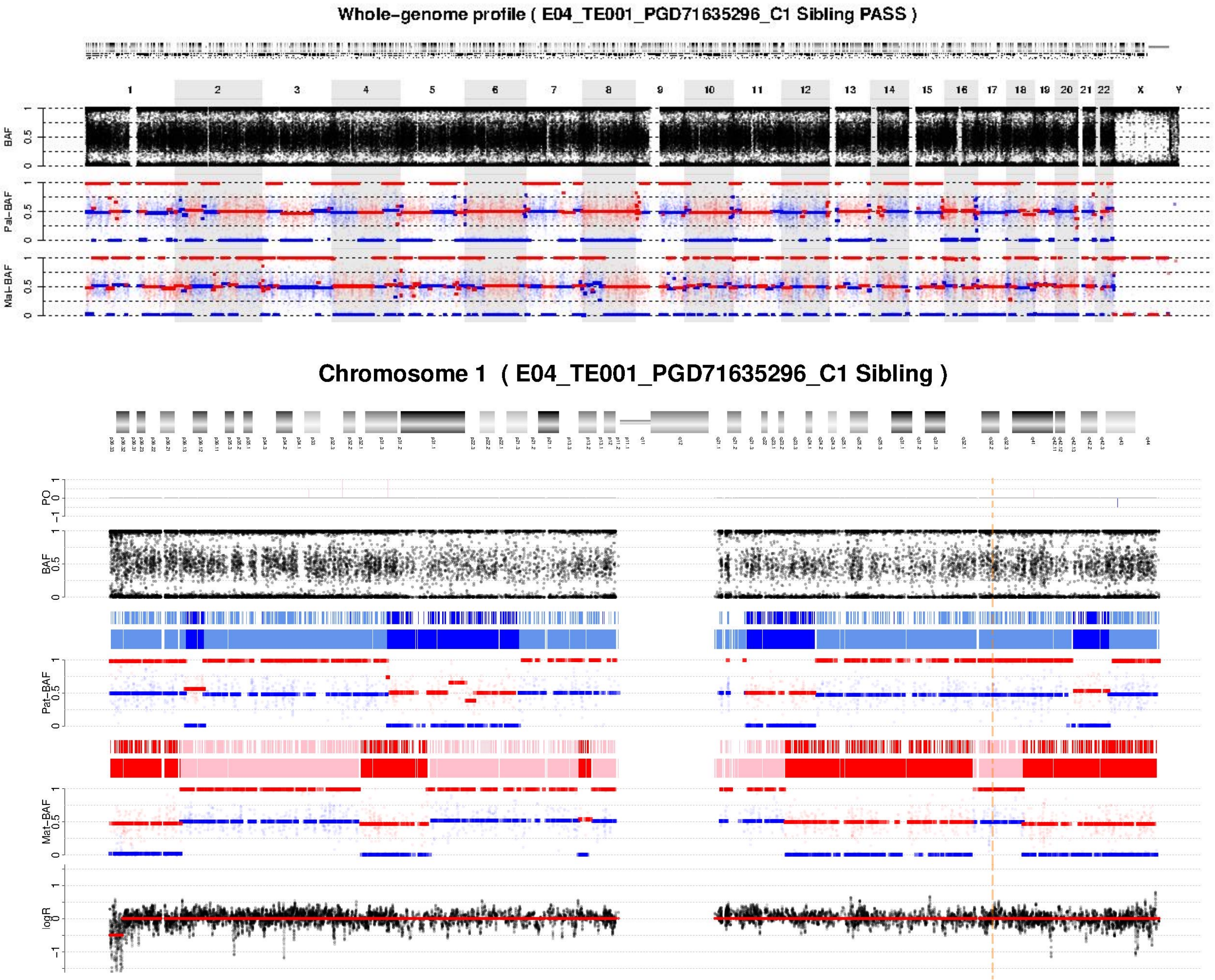


PGT-M:
Paternal SNV in *LAMB3*: not inherited
Maternal indel in *LAMB3*: not inherited

PGT-A:
Euploid

**D** ONT2-E06

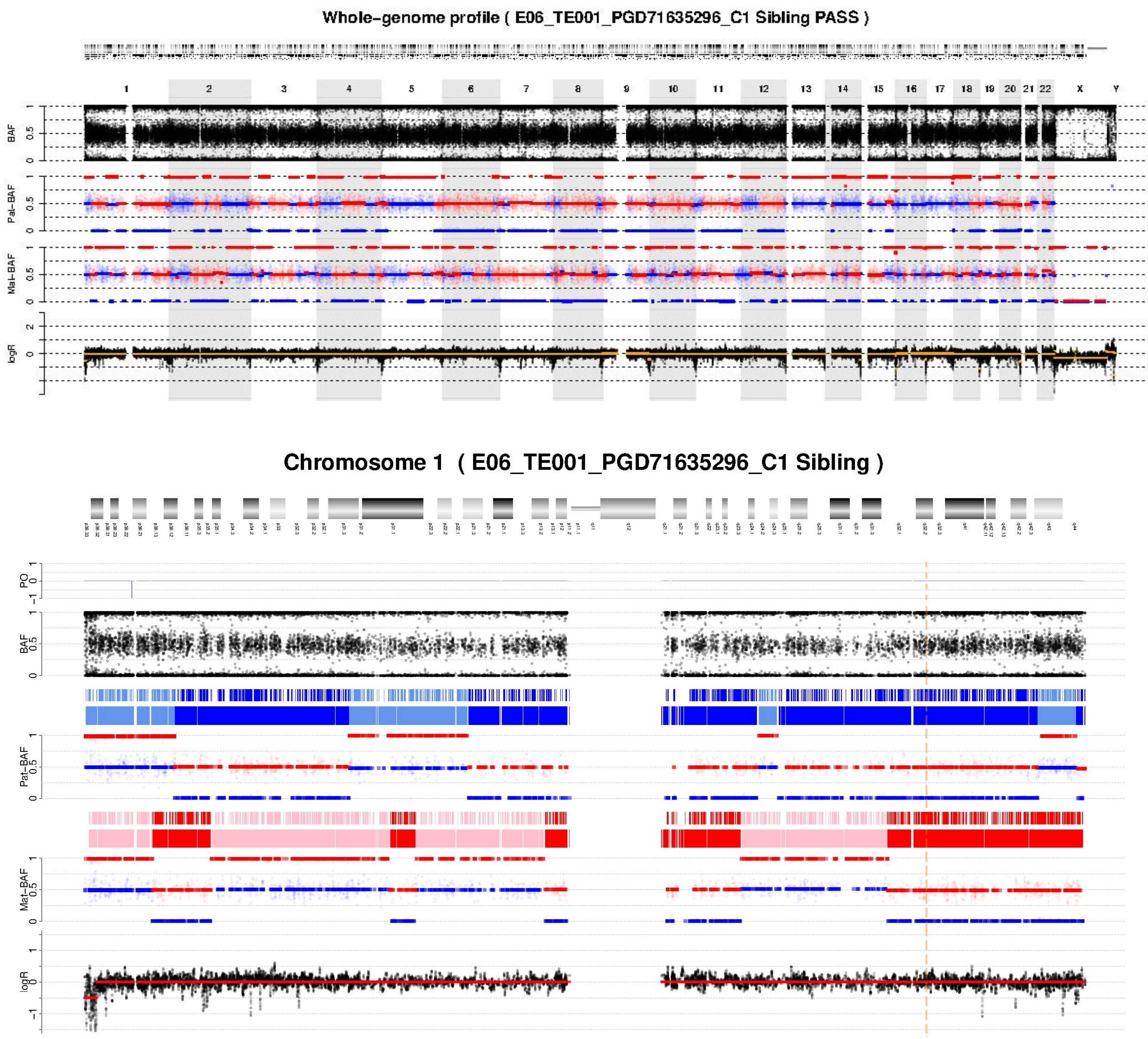


PGT-M:
Paternal SNV in *LAMB3*: inherited
Maternal indel in *LAMB3*: inherited

PGT-A:
Euploid

**E** ONT2-E20

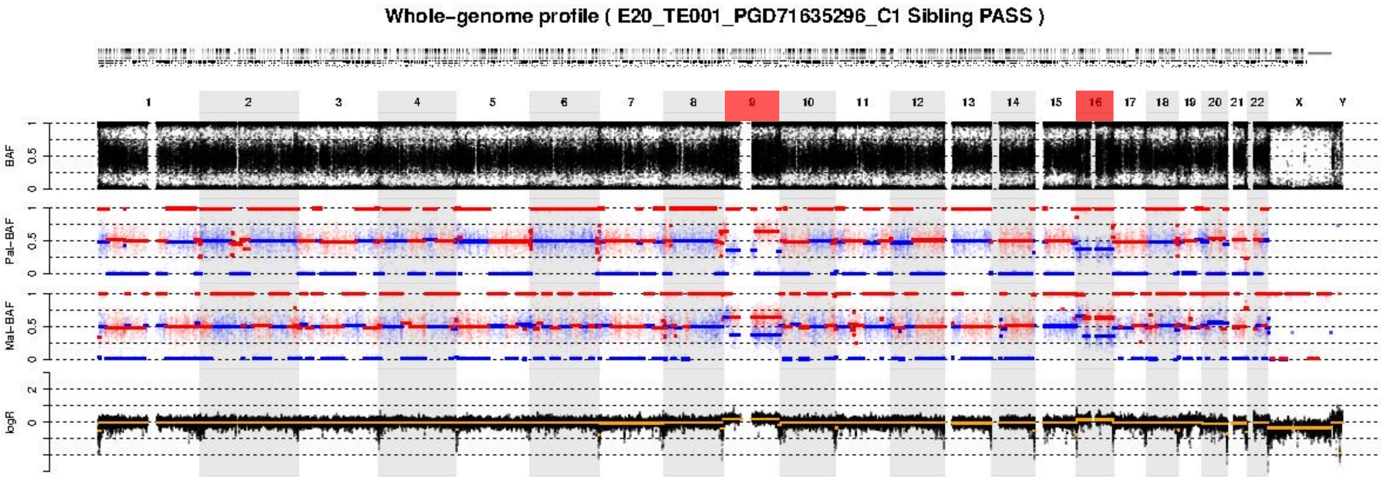


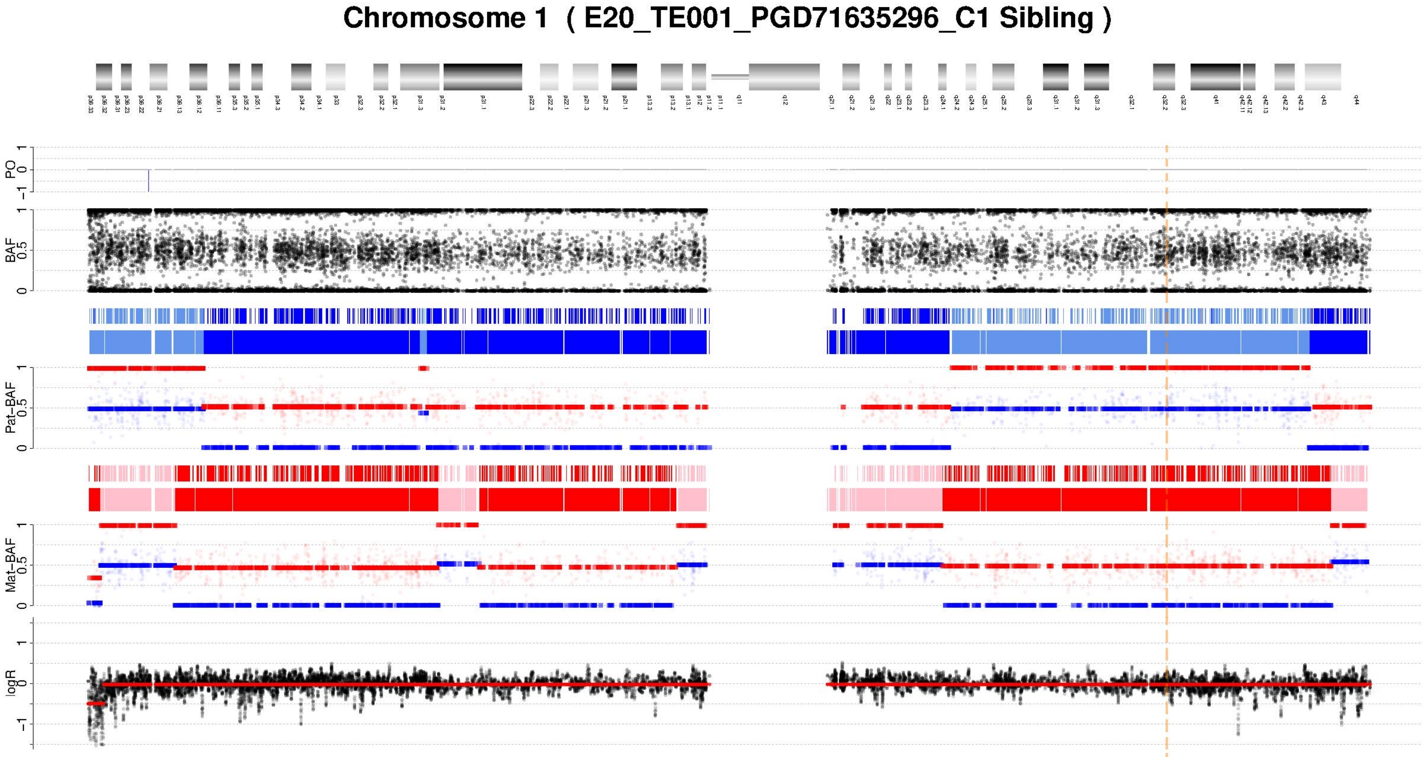


PGT-M:
Paternal SNV in *LAMB3*: not inherited
Maternal indel in *LAMB3*: inherited

PGT-A:
Trisomy for chromosomes 9 and 16, each involving an extra maternal chromosome resulting from nondisjunction during maternal meiosis I.

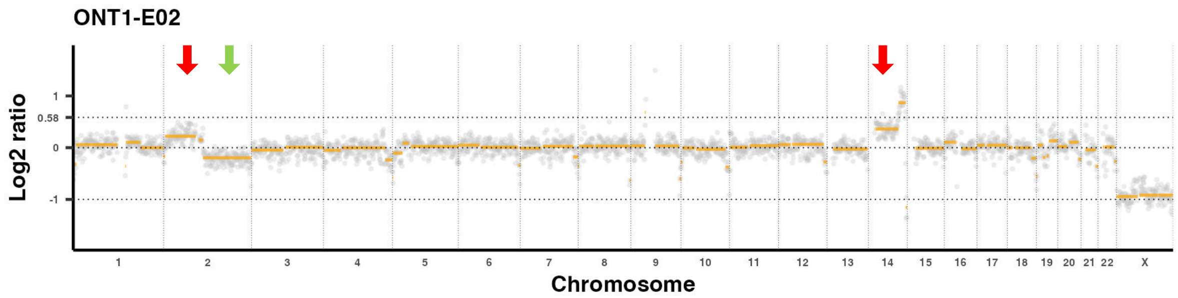


**Supplementary Figure 7.** PGT-A analysis using lrWGS data identified mosaic chromosomal abnormalities in ONT1-E02. The copy number plot reveals mosaic duplication of the short arm of chromosome 2 (indicated by a green arrow) and mosaic deletion of the long arm (indicated by a red arrow). Additionally, mosaic duplications are observed on chromosome 4 (indicated by a red arrow).
